# Molecular determinant of sterol recognition by sterol cleavage activatory protein

**DOI:** 10.1101/2024.11.19.624286

**Authors:** Charal Khiewdee, Puey Ounjai, Tanadet Pipatpolkai

**Author notes:** Corresponding author: *For correspondence, please contact:* Dr Tanadet Pipatpolkai, Dr Puey Ounjai.

## Abstract

Intracellular cholesterol homeostasis is regulated by the sterol response element-binding protein (SREBP) pathway through the SREBP cleavage-activating protein (SCAP) and the insulin-induced gene protein (INSIG). Previous studies highlighted that high concentrations of cholesterol mediated the dimerisation between INSIG and SCAP, which led to an inhibition of cholesterol uptake and biogenesis. However, the molecular understanding of SCAP-INSIG and cholesterol interactions remains elusive. Here, we used coarse-grained (CG) and atomistic (AT) molecular dynamics (MD) simulation to determine interactions between INSIG, SCAP and cholesterol. Our work highlighted the novel binding pocket in the luminal domain at the atomistic resolution. Using a combination of the AlphaFold3 model and Gō-martini forcefield, we showed that loop 7 dynamics are crucial for cholesterol binding and are able to highlight conserved residues, which lead to INSIG-SCAP dimerisation. Together, our work highlights novel mechanisms of cholesterol sensor pathways and will benefit the development of novel therapeutic strategies for diseases, such as neuroinflammatory disease, caused by irregular cholesterol homeostasis.

## Introduction

Cholesterol is one of the essential lipids constituting approximately 30% of the overall mammalian cell membrane, and it plays a significant role in regulating membrane rigidity, thickness, and permeability (*1–4*). In addition, cholesterol also serves as a precursor for steroid biosynthesis, such as progesterone and oestrogen (*5–7*). Dysregulation of cholesterol homeostasis, both in excess and in depletion, is a primitive cause of heart diseases and mental disorders, respectively (*8–10*).

The sterol regulatory element-binding protein (SREBP) pathway senses cholesterol levels in the endoplasmic reticulum (ER) (*11*). It has been proposed that there is a critical threshold level of cholesterol at approximately 5 mol% of the total ER membrane lipids. Within the ER, this pathway consists of three transmembrane proteins: SREBP-2, insulin-induced gene protein (INSIG), and SREBP cleavage-activating protein (SCAP) (*11, 12*). When the cholesterol level drops below the threshold, the SCAP-SREBP complex will be translocated to the Golgi apparatus, leading to an increase of cellular cholesterol synthesis (*6, 7, 13*). Together, the pathway has a negative feedback loop with cholesterol as a regulator (*7, 11, 12*).

Recent structural studies have shed light on the structure-function relationship of INSIG and SCAP (*12*). Structurally, INSIG comprises 6 transmembrane helices; SREBP comprises 2 transmembrane (TM) helices and 2 cytosolic domains, a regulatory element (Reg), and a basic helix-loop-helix-leucine zipper (bHLH-Zipper); and SCAP comprises of 8 transmembrane helices, 2 luminal loops, 1 cytosolic loop, and a cytosolic domain (*11*). The TM2-6 of SCAP has been previously described as a sterol sensing domain (SSD) that can sense membrane cholesterol levels (*14*). Interestingly, SCAP possesses two extended loops in the lumen between TM1-2 (loop 1) and between TM7-8 (loop 7) (*11, 15*). The cryo-EM structure of INSIG-SCAP dimer has been solved together with steroid ring-containing molecules, oxysterol, and digitonin. Remarkably, both ligands were found occupying the same site between INSIG and SCAP (*16, 17*). These ligands are also recognised as facilitators of INSIG-SCAP dimerisation by inducing INSIG and acting like a complex glue (*17*). However, the lack of experimental studies on this specific binding site hinders the results of the structural studies, leading to a contradiction in the current complex-glue hypothesis.

Besides, the previous study of a chicken SCAP (cSCAP) has hypothesised that the cholesterol binding site is potentially located on the luminal domain with an association of luminal loops 1 and 7 (*18*). This is supported by the change in the chicken SCAP conformation, especially in the TM7&8. The study has shown that the structure of cSCAP with a point mutation at D435V resembles the structure of human SCAP (hSCAP) of the 25-hydroxycholesterol-activated INSIG-SCAP complex (*18*). Remarkably, the comparison of cSCAP and cSCAP^D435V^ shows changes in TM7-8 locations after the mutation, leading to the change in luminal loops 1 and 7 (*18*). This study convinces that the interaction of cholesterol that induces the INSIG-SCAP dimerisation may potentially affect the luminal domain of SCAP (*16, 18*). However, the structure of cholesterol binding on SCAP has not been solved both in the TM and LM domains.

Here, our work uses coarse-grained (CG) and all-atom (AT) molecular dynamics (MD) simulations and reveals a low affinity of cholesterol interaction on the TM domain of INSIG and SCAP, which rejects the cholesterol complex glue in the INSIG-SCAP complex. We also report the calculated criteria from PyLipID used for the cholesterol-binding site identification. Our data emphasises that residues on luminal loops 1 and 7, which are highly conserved amongst animals, are important for cholesterol recognition and the subsequent conformational change. Together, our work helps clarify the key location of the cholesterol-binding on SCAP and generates a promising hypothesis on its cholesterol recognition, which can have implications for developing therapeutics for cholesterol-related diseases.

## Materials and methods

### Structural model and CG simulation preparation

The transmembrane segment (TM) of INSIG (residue 18-54/61-175/185-214) and SCAP (residue 5-47, 282-450, 535-560) were extracted from the structure from the Protein Data Bank (PDB ID:6M49) (*17*). The SCAP structure with a luminal (LM) domain (residues 10-49/69-450/625-661) was extracted from another INSIG-SCAP complex (PDB ID:7ETW) (*16*). In the segment of the work where the role of loops 1 and 7 are studied, loop 1 (residues 90 to 280) or loop 7 (residues 625 to 661) were removed from the structure. Each structure was converted into coarse-grained (CG) using *martinize2* with either MARTINI2.3, MARTINI3 or Gō-MARTINI forcefield (Table 2) (*19–21*). with an elastic network with a force constant of 1000 kJ/mol/nm^2^ between backbone beads within 0.6–1.0 nm to maintain the protein structure. In the simulations where we are interested in the dimerisation interfaces, the TM-INSIG and TM-SCAP were placed 50 Å away from each other. The protein(s) CG structures were then embedded individually in DLPC/cholesterol lipid bilayers to mimic an ER membrane environment using *insane.py* (*22, 23*). The ratio of DLPC/cholesterol is different depending on each investigation, as denoted in Table 1. All systems were then flooded with coarse-grained water particles with 0.15 M NaCl (*24*). The protein(s) were placed away from the edge of a box about 20 Å. All structures were visualised using PyMOL (*25*).

**Table 1.** The details of the investigations and the membrane composition setup.

| Investigation aims | The structures | Membrane compositions |
| --- | --- | --- |
| To observe the cholesterol interaction on TM domains | TM-INSIG & TM-SCAP | 85% DLPC: 15% cholesterol |
| To observe the cholesterol-inducing INSIG-SCAP dimerisation | TM-complex-M2 & TM-complex-M3 | 85% DLPC: 15% cholesterol |
| To investigate an effect of cholesterol on luminal domain dynamics | SCAP with Gō-martini | 70% DLPC: 30% cholesterol |
| To observe the stability of cholesterol within the INSIG-SCAP complex cleft | TM-cleft series | 100% DLPC |
| To observe the cholesterol interactions on the luminal domain of SCAP | SCAPΔL7 & SCAP | 100% DLPC |

**Table 2.** The overview of all simulations.

| **Systems** | **Duration (μs)** | **Total (us)** |
| --- | --- | --- |
| TM-INSIG (CG) | 5.5 x 5 | 27.5 |
| TM-SCAP (CG) | 5.5 x 5 | 27.5 |
| TM-complex-M2 (CG) | 20.2 x 3 | 60.6 |
| TM-complex-M3 (CG) | 5.2 x 3 | 15.6 |
| TM-cleft (CG) | 5.2 x 3 | 15.6 |
| TM-cleft-CHOL (AT) | 0.25 x 3 | 0.75 |
| TM-cleft-25HC (AT) | 0.25 x 3 | 0.75 |
| SCAPΔL7 (CG) | 0.2 x 100 | 20 |
| SCAP (CG) | 0.2 x 100 | 20 |
| SCAP (Gō-martini) with<br>CHOL | 5x 5 | 25 |
| SCAP (Gō-martini) without<br>CHOL | 5x 5 | 25 |
| SCAP-APO (AT) | 0.5 x 3 | 1.5 |
| SCAP-CHOL (AT) | 0.5 x 3 | 1.5 |

### Cholesterol interaction with SCAP and INSIG

Coarse-grained cholesterol molecules were placed between SCAP and INSIG, randomly in the membrane, or randomly replaced water particles with 10 cholesterol molecules using *gmx insert-molecule* (*26*). All systems were energy minimised using a steepest-descent algorithm for 5000 steps with an energy cap of 1000 kJ mol^-1^ nm^-2^ (*27*). The length and the equilibration time of all simulations are described in Table 2. All CG-MD simulations were performed with 20 fs timesteps. All CG-MD simulations were performed using GROMACS 2021.3 (*26*). The temperature was maintained at 310 K using V-rescale thermostat temperature coupling (*28*). The Parrinello-Rahman barostat was used to control semi-isotropic pressure at 1 bar in xy direction (*28, 29*). All minimum distance analyses were performed using GROMACS 2021.3 (*26*). The cholesterol binding site was then calculated using the PyLipID double cutoff scheme at 0.5 and 0.8 nm (*30*).

### Coarse-grained-free energy perturbation (CG-FEP)

The cholesterol-bound systems were obtained from the last frame snapshot of each CG-MD simulation. The systems were edited to remove other cholesterol molecules, leaving only one molecule at the binding site using PyMOL (*25*). The free energy perturbation (FEP) method calculates binding free energy (ΔΔG) by converting a coarse-grained cholesterol molecule into a dummy molecule with no electrostatic or Lennard-Jones interaction potential along the λ chemical space coordinate from the step of λ = 0 (cholesterol) to λ = 1 (dummy molecule) using 10 windows (λ = 0.00, 0.10, 0.20,…, 1.00) (*31*). The FEP was performed in both bound states (with protein) and a solution box with CG water particles and 0.15 M NaCl, creating a thermodynamic cycle of cholesterol binding (*24*). The transformation was conducted with the soft-core parameters of α = 0.5 and σ = 0.3. In every window, the system was energy minimised with the steepest descent algorithm for 500 steps. Each FEP system was simulated using a leap-frog stochastic dynamics integrator to 12 ns per window. The first two ns were discarded as an equilibration for 15 repeats per each condition. The Multistate Bennett Acceptance Ratio (MBAR) was used to calculate the binding free energy of each simulation window and, thus, the overall binding free energy (*32*). All calculations were conducted using *alchemical-analysis.py* Python packages (*33*).

### All-atoms molecular dynamics simulation

In the TM-cleft-CHOL&25HC systems, the INSIG and SCAP structures were placed in the DLPC bilayers using the CHARMM-GUI Membrane Input Generator (*34*). Afterwards, the generated ligands were placed in the same pose as in the experimental results (PDB ID: 7ETW and 6M49) (*16, 17*). Another, the equilibrated CG-SCAP systems were converted into all-atom (AT) using CG2AT2 (*35*). The topology structures of cholesterol analogues were then added. The topology of cholesterol (SCAP-CHOL) and 25-hydroxycholesterol (25HC) were generated using the CHARMM-GUI Ligand Input Generator (*36*). All ligands were placed in the cholesterol binding site. In the all-atoms configuration, the CHARMM36m forcefield with TIP3P water model was used throughout this study (*37*). The systems were then energy minimised using a steepest-descent algorithm for 5000 steps and equilibrated for 10 ns with the restraint of 1000 kJ nm^-2^ mol^-1^ on C_α_ of the protein and every atom of the cholesterol molecule (*27*). The systems were then equilibrated for 10 ns, maintaining the temperature at 310 K using a V-rescale thermostat and the Berendsen barostat for semi-isotropic pressure coupling at 1 bar (*28, 38*). Then, the system was simulated following Table 2. The AT-MD simulations were simulated using 2 fs timesteps and maintained the temperature at 310 K using a V-rescale thermostat and the Parrinello-Rahman barostat for semi-isotropic pressure coupling at 1 bar (*28, 29*). All simulations and analyses were performed using GROMACS 2021.3 (*26*).

### Multiple sequences alignment (MSA)

The human SCAP (Uniprot Q12770, residues 1-1279), chicken SCAP (Uniprot A0A3Q3ANV4, residues 1-1316), mouse SCAP (Uniprot Q6GQT6, residues 1-1276), frog SCAP (Uniprot A0JPH4, residues 1-1311), fruit fly SCAP (Uniprot A1Z6J2, residues 1-1276), worm SCAP (Uniprot Q18968, residues 1-1087) and yeast SCAP (Uniprot O43043, residues 1-1086) were aligned and visualised with MultAlin (*39, 40*). The phylogenetic tree was constructed from the Clustal Omega web server using the multiple sequence comparison by log-expectation (MUSCLE) algorithm (*41*).

### AlphaFold and Chai-1

The amino acid sequence of human SCAP (Uniprot Q12770, residues 1-1279) and INSIG were inputted to the Google DeepMind AlphaFold3 to predict the binding interface between SCAP and INSIG and or their cholesterol binding site (*42*). The model with the highest ranked score was chosen for cholesterol docking using Chai-1 (*43*). The 9 cholesterol molecules were input together with SCAP and SCAP dimerising with INSIG, generated from Alphafold3. The results were visualised using PyMOL (*25*).

## Results

### The cholesterol does not bind to the transmembrane of SCAP stably

The dimerisation event between INSIG and SCAP is induced by sterols (*11*). Recent cryo-EM structures of INSIG-SCAP (PDB entry: 6M49 & 7ETW) show steroid rings (25-hydroxycholesterol and digitonin) between INSIG and SCAP in the complex (*16, 17*). However, previous studies showed that loop 1 and loop 7 are important in the cholesterol sensing processes, and thus, whether cholesterol is important for the dimerisation of INSIG and SCAP in the transmembrane (TM) domain marked a significant question (*18*). To answer, we conducted the CG-MD simulations of TM-INSIG and TM-SCAP within the excess-cholesterol DLPC membrane, 15% cholesterol for 5.5 us (n = 3) (Figure 1A, Table 1) (*14*). We hypothesised that cholesterol molecules potentially occupied the TM segments of INSIG and SCAP for a long period due to the very distinct cryo-EM density (*17*). Here, we calculated the minimum distance between cholesterol and TM-INSIG and TM-SCAP. Our simulations show that cholesterol molecules do not reside stably within the binding site solved from the cryo-EM structure (Figure 1B) (*16, 17*). To force cholesterol to bind stably in the binding pocket, we placed a cholesterol molecule in the middle of the INSIG-SCAP dimerisation interface and tracked an interaction between the ROH beads on the cholesterol molecules and K107 of INSIG and A132 of SCAP (Supplementary Figure 1A). We expected a stable binding without flipping of the ROH beads in the interface. However, the cholesterol molecule presents an unstable binding by rotating within the cleft in 3 out of 5 simulations (Supplementary Figure 1B). From this, our results highlight that the transmembrane regions of SCAP and INSIG are unlikely to be the cholesterol-binding sites.

**Figure 1.**
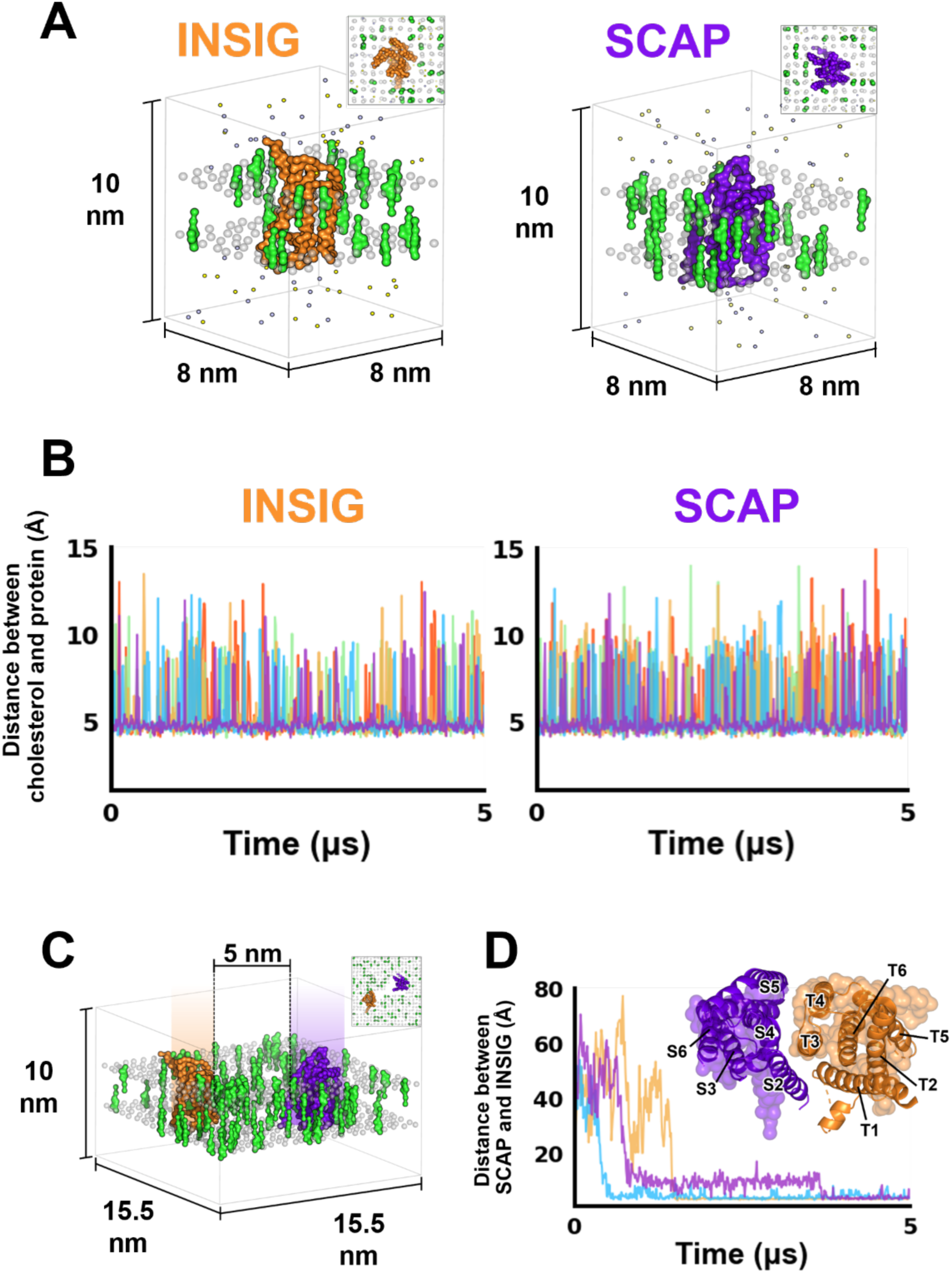
CG-MD simulations and the transient contacts of cholesterol on the transmembrane domain. **(A)** The overview of CG systems containing TM-INSIG (left) and TM-SCAP (right), which are embedded in the DLPC lipid bilayers (phosphate headgroups are shown in grey) with cholesterol (green). Water (not shown) and ions are shown as tiny spheres. The top views of each box are shown in the upper-right corner of each system. **(B)** The graph shows the distance between cholesterol and the TM-INSIG (left) and TM-SCAP (right). Each colour indicates different repeats (n=5). **(C)** The overview of the CG system containing TM-complex, TM-INSIG (orange) and TM-SCAP (purple), which are placed away from another *ca.* 5 nm and embedded in the same building as (A). **(D)** The graph shows the distance between TM-INSIG and TM-SCAP within the TM-complex systems. Each colour indicates different repeats (n=3). The configuration that matches the experimentally solved structure of the INSIG-SCAP complex is shown as translucent molecules at the upper-right corner of the figure, superimposed with the experimental result in cartoon.

Previous simulations show that cholesterol only binds to INSIG and SCAP monomers transiently (Figure 1B). We then asked whether cholesterol induces the dimerisation of INSIG and SCAP without binding to the molecules. We placed SCAP and INSIG 5 nm away from each other in a 15×15×10 nm simulation box and conducted 20.5×3 µs simulations in a 15% cholesterol DLPC membrane (TM-complex) (Figure 1C). We hypothesised that cholesterol should mediate dimerisation and reside between SCAP and INSIG. Here, our results show that TM-INSIG and TM-SCAP dimerised each other without any cholesterol between them, highlighting that cholesterol is not required for the SCAP-INSIG dimerisation in the transmembrane domain (Figure 1D). In addition, we suspect that the lack of observable dimerisation is due to the coarse-grained forcefield. Thus, we conducted the simulations in both the MARTINI2 and MARTINI3 forcefield and yet, we still do not observe cholesterol binding at the dimerisation interface (Supplementary Figure 2), despite getting the correct protein interaction interface from the MARTINI3 forcefield (Figure 1D). In addition, Together, our work highlighted that the interaction of cholesterol in the transmembrane region does not influence the dimerisation between the transmembrane region of the two proteins - contradicting the structural data.

### 25-HC binds to SCAP and INSIG in the transmembrane region

We asked the question on the role of cholesterol and 25-HC in the transmembrane region given that the structure has been previously solved with 25-HC and digitonin at the binding cleft in the transmembrane region (*16, 17*). Previous studies have shown that 25-HC binds to SCAP at the transmembrane region, but cholesterol does not (*17*). To gain insight into this observation, we conducted 500 ns atomistic simulations of SCAP-INSIG dimer, with 25-HC or cholesterol in the binding interface between two proteins. Here, we show that with 25-HC, the hydroxyl at the tail forms a hydrogen bond with T147 on INSIG and stabilises the tail conformation (Figure 2A). On the other hand, in the conformation with cholesterol at the dimer interface, the tail shows fewer stable dynamics, leading to a slightly higher root mean square deviation (RMSD) (Figure 2B). This is in agreement with our hydrogen bond analysis, where 25-HC forms more hydrogen bonds (2 to 3 H-bonds) with the protein. In contrast, cholesterol primarily only forms a single hydrogen bond with SCAP. Together, our results explain the molecular basis of the 25-HC binding within the transmembrane domain and explain the lack of stability of cholesterol binding in the transmembrane region.

**Figure 2.**
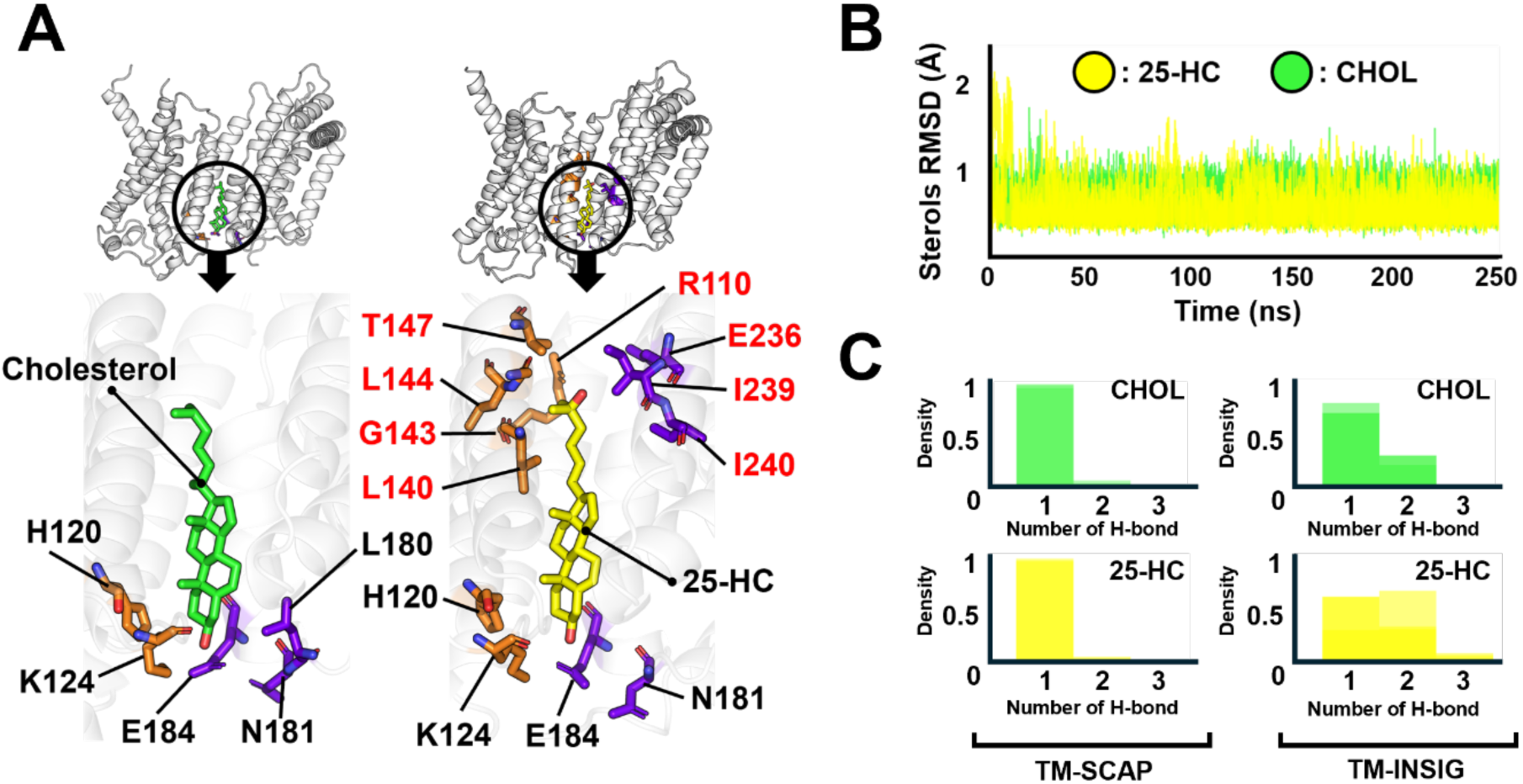
The difference between cholesterol and 25-hydroxycholesterol (25-HC) in binding within a cleft of TM-complex. **(A)** The PyLipID generated a binding site for cholesterol (green) and 25-HC (yellow). Each contributing residue is calculated by PyLipID. Residues those are only contacting with 25-HC are shown in red. The residues of TM-INSIG are shown in orange, and TM-SCAP are shown in purple. **(B)** The RMSD values of the cholesterol (green) and 25-HC (yellow) molecules throughout 250 ns simulation (n=3). **(C)** The distribution of H-bonds between ligands, cholesterol (green) and 25-HC (yellow), and TM-SCAP (left) and TM-INSIG (right) (n=3 of each).

### The cholesterol binding site is located in the luminal domain of SCAP

Given that our previous simulations only reveal a transient interaction of cholesterol on the TM domain and fail to demonstrate that cholesterol is required for the dimerisation of INSIG and SCAP, we change our attention to the luminal domain (LM), the region which has been previously experimentally shown to be related with cholesterol sensing function (*18, 44*). However, there has been no structural or computational evidence highlighting the importance of loop 1 and loop 7 on the cholesterol sensing mechanism. To computationally look for the possibility of cholesterol binding to loop 1 and loop 7, we used PocketMiner to identify the cryptic pocket within the SCAP structure based on the consensus topology and sequences within the database (*45*). The result highlights 2 potential regions on the sterol-sensing domain (SSD), which is located at the transmembrane region (residue 284-442) and within the luminal domain of SCAP (Figure 3A-B). This convinces us that the cholesterol-binding site is located in the LM domain. We then conducted 100 replicates of 200 ns CG-MD simulations. Each system contains 10 cholesterol molecules randomly placed in the solvent (Figure 3C). We hypothesised that the cholesterol molecule would enter the potential cholesterol binding region on the LM domain. Here, we reveal one cholesterol binding site where a cholesterol molecule occupies more than ∼40% of the simulation time of 100 repeats, matching with the site identified by PocketMiner. We define occupancy as the cholesterol molecule remaining 0.5-0.8 nm within the binding pocket using PyLipID (Supplementary Figure 3). From 37 out of 100 repeats, cholesterol entered and occupied the identified cholesterol binding site, i.e. cholesterol binding site 7. The other, at 63 repeats, cholesterols migrate towards the phospholipid bilayer, leading to no interaction with the SCAP molecules. This cholesterol binding site is located between loop 1 and loop 7 of SCAP (Figure 3D).

**Figure 3.**
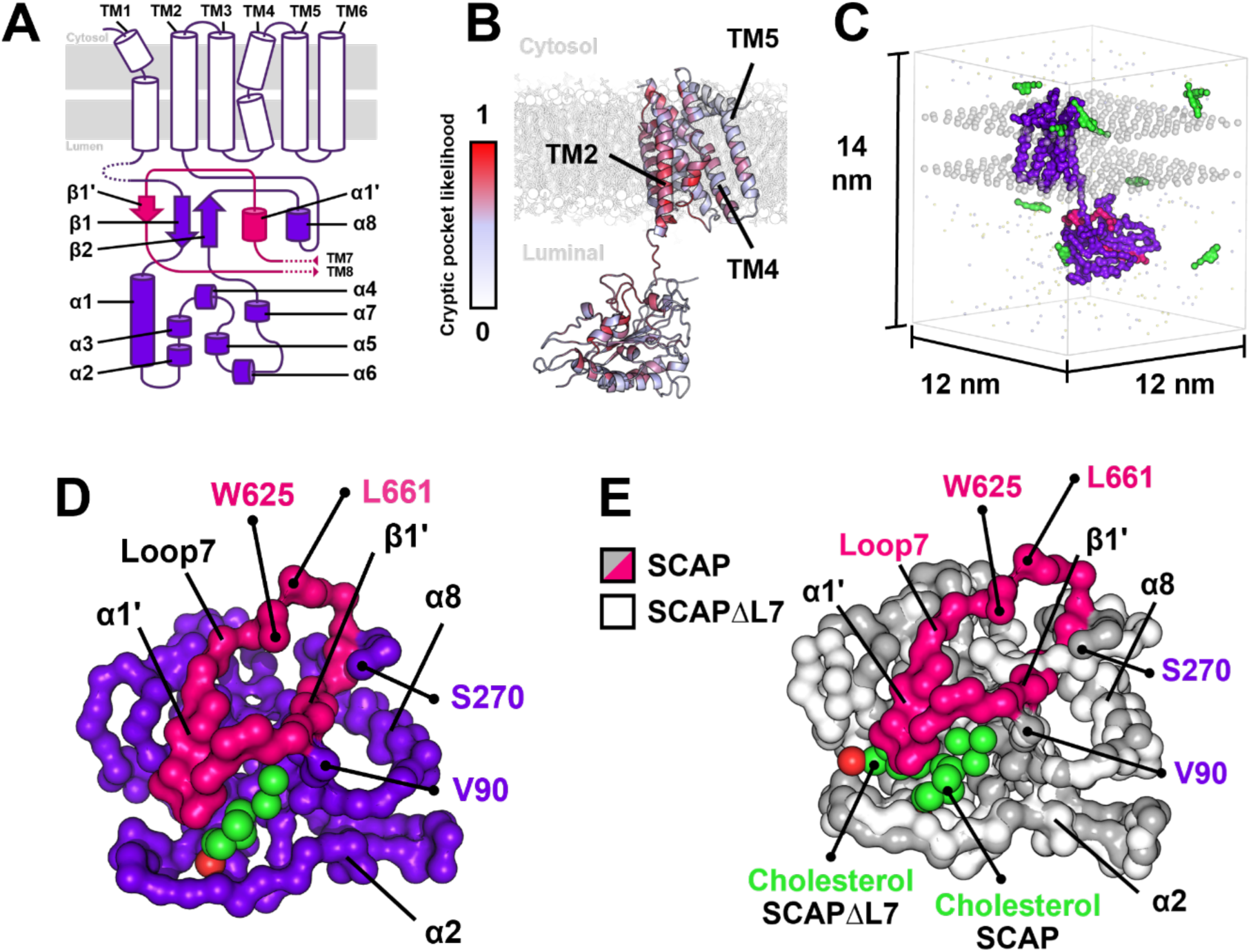
The interaction of cholesterol on the luminal domain of SCAP. **(A)** The topology of the SCAP structure that is used in this study shows the transmembrane domain (white), luminal loop 1 (purple), and luminal loop 7 (hot pink). **(B)** The structure of SCAP shows different colours in each part, which indicate different potential cryptic pocket scores generated by Pocket Miner. The score colour reference is shown on the left side; 1 is the highest score, and 0 is the lowest score. **(C)** The overview of CG systems containing SCAP which is embedded in the DLPC lipid bilayers (phosphate headgroups are shown in grey). 10 added CG-cholesterol molecules are shown in green. Water (not shown) and ions are shown as tiny spheres. The TMs and luminal loop 1 are shown in purple, while the luminal loop 7 is shown in hot pink. **(D)** The LM domain of SCAP shows the cholesterol (green) binding area on the luminal loop 1 (purple) and 7 (hot pink). **(E)** The superimposition of LM domain of SCAP and SCAPΔL7 shows an overlapping area of cholesterol (green) binding.

Given that both loops are required for cholesterol interaction, we were interested in the requirement of each loop in the cholesterol recognition mechanism. To do so, we truncated loop 7 (SCAPΔL7) and conducted the same simulation as above. Surprisingly, using similar analyses, we show that the binding site remains unchanged, likely due to a very subtle effect (Figure 3E, Supplementary Figure 3). To address such a subtle difference, we used CG-FEP (coarse-grained free energy perturbation) to calculate the affinity of the cholesterol molecule to the binding site. We converted a cholesterol molecule to a dummy molecule and calculated free energy differences in steps (ΔΔG) (Supplementary Figure 5). The results reveal significantly more negative binding energy when both loops are presented (ΔΔG = −47.06 kJ/mol) compared to the protein without loop 7 (ΔΔG = −42.29 kJ/mol) (Supplementary Figure 4). From this, our results highlight that the cholesterol binding site is located between loop 1 and loop 7 of SCAP, with loop 7 potentially stabilising the cholesterol binding site.

We then aim to gain an atomistic insights towards the cholesterol binding pocket. Thus, we turn to an all-atoms (AT) simulation of SCAP, where we simulated SCAP with cholesterol in the binding pocket for 500 ns (n=3) to observe the dynamic of the cholesterol molecule within the binding pocket. The PyLipID analysis reveals a group of amino acids occupied by the cholesterol molecule for more than 95% of the simulation time, with 0.5-0.8 nm dual cut-off. These amino acids were then characterised as the cholesterol binding site of our study (Figure 4A). In our studies, all binding modes defined from PyLipID are relatively similar, with loop 1 interacting with the hydroxyl group and loop 7 stabilising the tail of the cholesterol molecule (Supplementary figure 5) (*30*). Within this binding site, the cholesterol molecule is primarily making contact with V93, L148, L164, L170, L172 and V239 on loop 1 and F637, Y640, I642 and I649 on loop 7 (Figure 4B). Interestingly, the hydroxyl group of the cholesterol molecule has been shown to interact with the backbone amide of L148, given that there is no nearby polar side chain. This binding mode is stable, given the RMSD of the ligand is within 3 Å of the initial configuration (Figure 4C).

**Figure 4.**
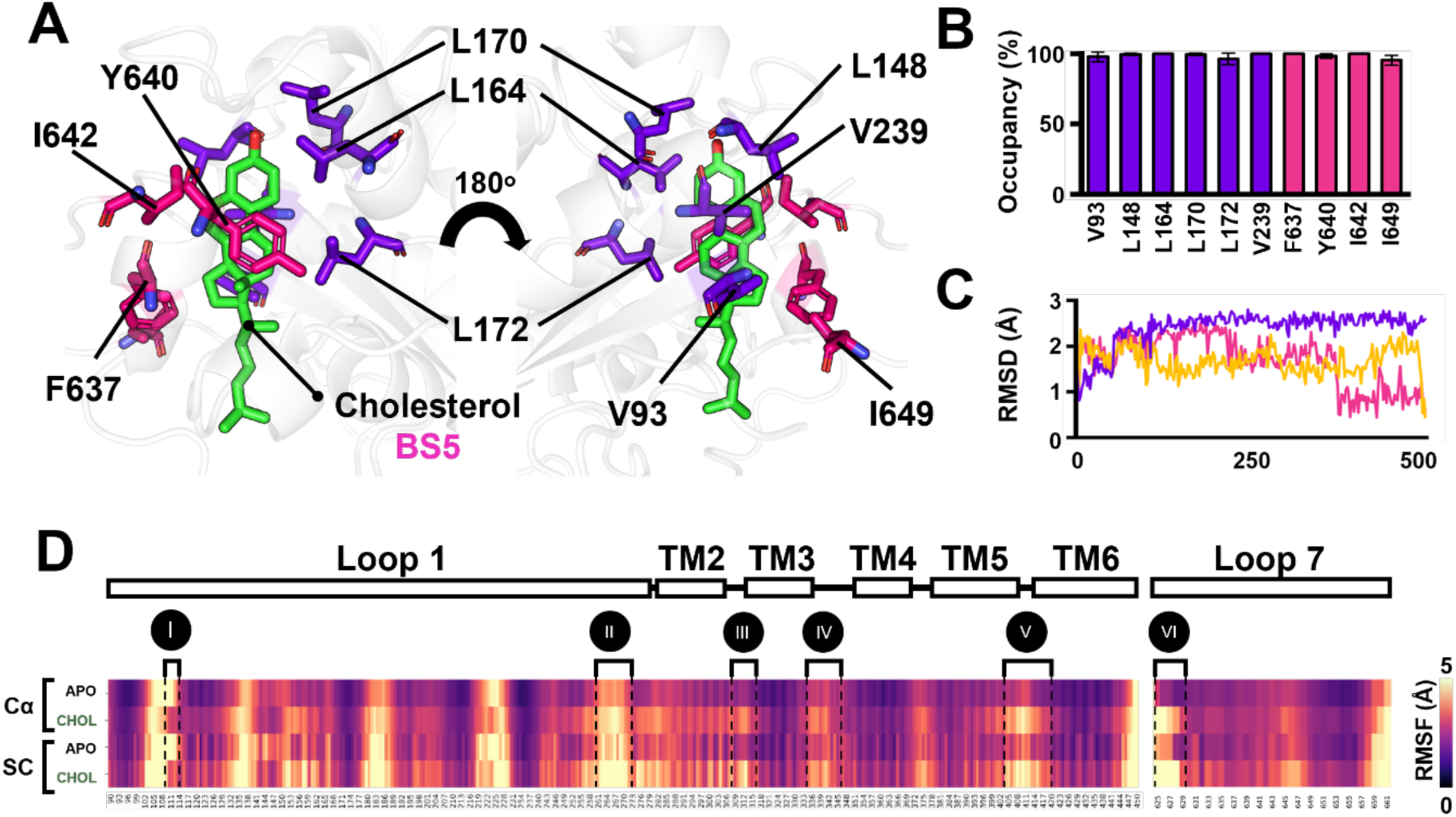
The atomistic detail of cholesterol binding side. **(A)** The cholesterol binding site shows selected residues that interact with a cholesterol molecule (green). The residues were selected according to the highest score in duration, occupancy, lipid counting, and residence time criteria (Supplementary figure 5A-E). The cholesterol pose was obtained from BS5. The residues of luminal loops 1 and 7 are shown in purple and hot pink, respectively. **(B)** The graph of selected residues of luminal loop 1 (purple) and 7 (hot pink) from (A) shows >95% cholesterol occupancy scores throughout the 500 ns (n=3). **(C)** The RMSD analysis of cholesterol within SCAP-CHOL system shows changes in cholesterol pose compared with the cholesterol pose of the first frame (t=0). The different colours represent different repeats (n=3). **(D)** The RMSF results show the fluctuation of Cα and side chain (SC) of SCAP-APO and SCAP-CHOL. Secondary structural elements of TM (residues 281-450) and LM (residues 90-280/625-661) domains of SCAP are shown above the RMSF results. The regions that present an obvious change in RMSF are shown in the different Roman numbers, I-VI.

Next, we conducted root mean square fluctuation (RMSF) calculation on the residues within the cholesterol binding pocket in the presence and absence of cholesterol. Here, we showed that cholesterol binding stabilises the C_α_ and side chains regions I (residue 108-114) on loop 1 and destabilises region II (residue 261-2404-418 73), III (residue 308-315), IV (residue 336-342), V (residue 404-418), and VI (residue 625-630). These regions are very close to the cholesterol-binding pocket on loop 7, suggesting that the binding of cholesterol may indeed relay a conformational change down the loop and, thus, increase the dynamic property of loop 7. In addition, using multiple sequence alignment (MSA), we show that the region at which the RMSF is affected by cholesterol binding, both on loop 1 and loop 7, is highly conserved across the animal kingdom (Supplementary Figure 6). Together, our results reveal that all regions that present a high RMSF after the cholesterol binding are conserved in all selected organisms and thus may be involved in some crucial conformational change regarding cholesterol interaction.

### High cholesterol in the ER causes a conformational change in the luminal domain

In this study, we hypothesised that a cholesterol binding event in the luminal domain will trigger the conformational change, which leads to the dimerisation between SCAP and INSIG. However, cholesterol is a hydrophobic molecule and, thus, is unlikely to be in the lumen for the interaction at the luminal domain. By closely exploring the structure of the related protein, NPC1, we observed a putative tunnel where cholesterol could translocate from the membrane to the binding site of interest (Figure 5A, Supplementary Figure 7) (*46–48*). The structure of the C domain on the NPC1 is very similar to the luminal domain of SCAP, with the difference being the additional loop 7 blocking the translocation path (Supplementary Figure 7B) (*16, 47*). In addition, we use the Chia-1 AI generative model to predict the cholesterol binding site, and this shows that cholesterol does not reside at the dimerisation interface but is filling the cavity near TM7, an aforementioned putative tunnel (Supplementary figure 7) (*43*). Thus, we hypothesise that there must be a conformational change affected by cholesterol, allowing cholesterol to move to our luminal domain of interest for a large-scale conformational change. To address this, we conducted additional coarse-grained simulation with the Gō-martini forcefield, allowing the conformational change within the luminal domain to be observed more reliably than the traditional MARTINI-forcefield simulation system (*21*). We compare the dynamical motion of the luminal domain in the membrane with and without 30% cholesterol and measure the distances between the luminal domain and the phosphate headgroup in the bilayer over 5 µs for 5 repeats (Figure 5B). Our simulation shows that with the presence of cholesterol, the luminal domain is more in contact with the phospholipid bilayer (Figure 5C). Despite our limitation of not observing direct cholesterol translocation to the binding site, this suggested a hypothesis that increases in cholesterol concentration in the membrane lead to the conformational change down the luminal domain. We then used AlphaFold3 to predict the structure of SCAP in the absence of cholesterol (*42*). Here, the model suggested similar results to the previous experimental studies where cholesterol sensing leads to a large-scale conformational change (Supplementary Figures 8&9) (*18*). Without cholesterol, the AlphaFold3 models show that TM7 and TM8 block the dimerisation interface between SCAP and INSIG (Supplementary Figure 9). Thus, only when cholesterol is in excess in the membrane is the SCAP luminal domain translocated upward to the membrane, allowing the cholesterol to migrate down the tunnel to its binding site with the large-scale conformational change. This enables the interface between INSIG and SCAP to be revealed for INSIG-SCAP dimerisation.

**Figure 5.**
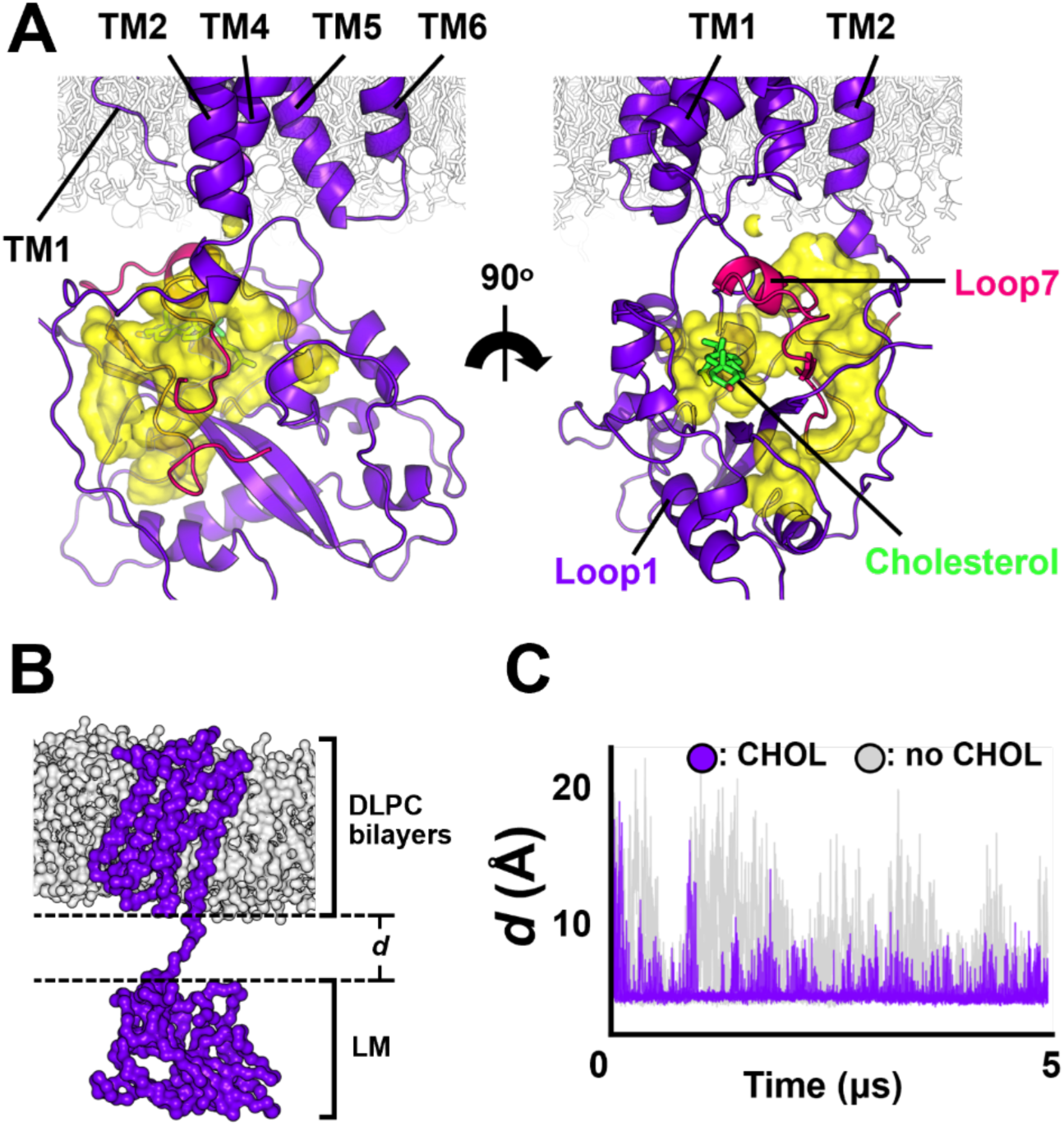
The potential conformational change of LM domain of SCAP. **(A)** The structure of LM domain of SCAP shows cavity (yellow) underneath the INSIG-dimerising interface (TM2/4/5) lining to the cholesterol (green) binding area on the luminal loops 1 (purple) and 7 (hot pink). **(B)** The overview of the initial state of CG-SCAP structure shows the minimum distance (*d*) between TM (residues 10-47/281-450) and LM domains (residues 90-275). The structure is embedded in the DLPC lipids bilayers (grey) containing 15% cholesterol (not shown). **(C)** The changes of *d* throughout 5 µs simulations of SCAP with cholesterol (purple) and with no cholesterol (grey) systems (n=3 of each).

**Figure 6.**
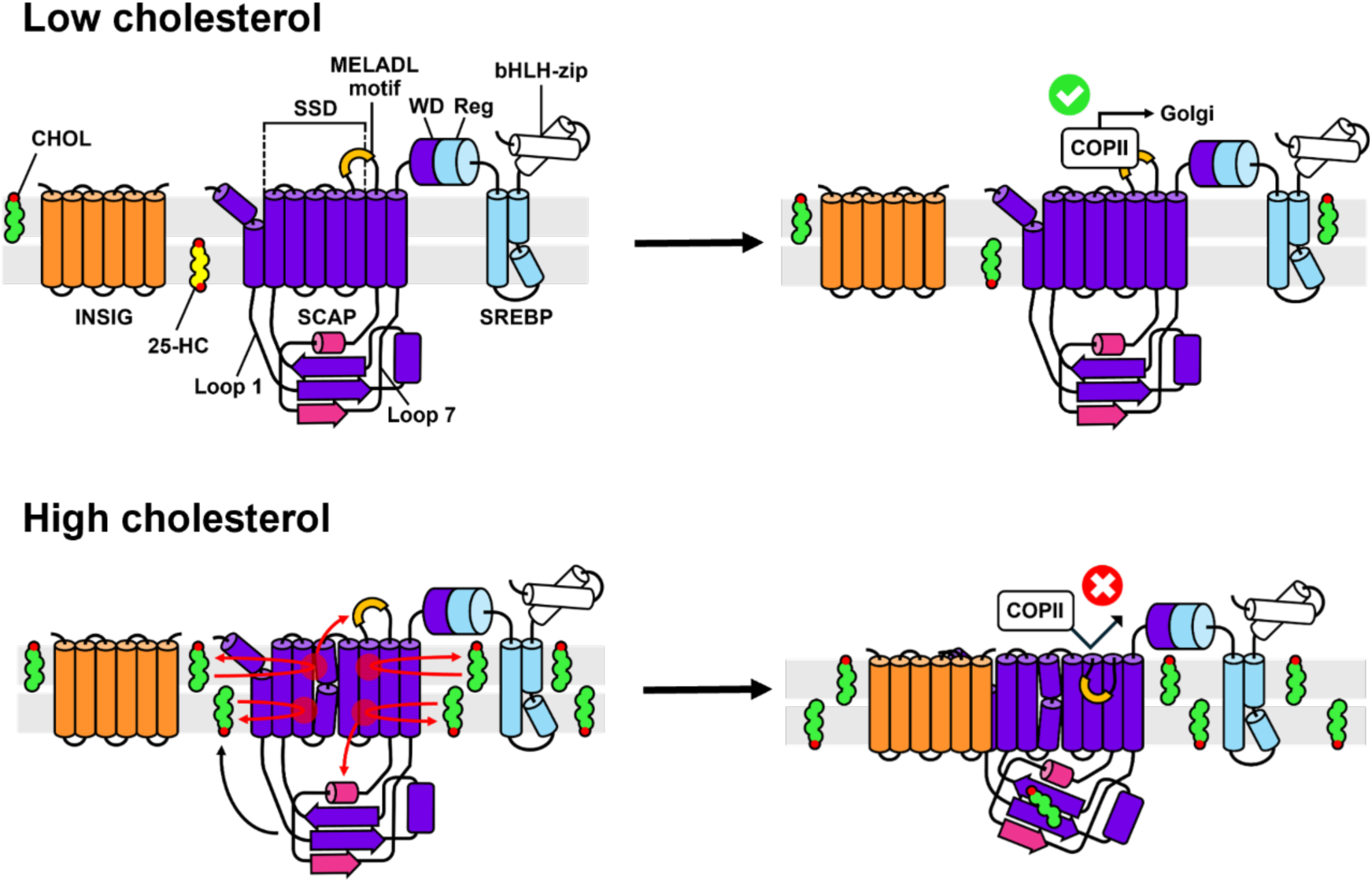
The schematic of SREBP pathway. (Upper panel) When the cholesterol level within the ER membrane is below 5 mol% of total ER lipids, COPII proteins enter and recognise the MELADL motif on the cytosolic loop of TM6 of SCAP (purple), allowing SREBP (light blue) to be translocated to the Golgi apparatus. (Lower panel) When the cholesterol level within the ER membrane is higher than 5 mol% of total ER lipids, the cholesterol molecules contact the SSD domain of SCAP more frequently, allowing the LM domain to move up and connect to the TM domain, allowing cholesterol to translocate to the cholesterol binding site on LM domain and hide the MELADL motif on TM6 consequently. This causes SCAP to dimerise with INSIG (orange). This causes an absence of the SCAP-SREBP complex translocation.

## Discussion

The SREBP pathway is inactivated by the interaction of cholesterol on SCAP (*11, 12*). Previous studies have reported alternative sterols that can inhibit the SREBP pathway by interacting with either INSIG or SCAP (*49*). An interesting alternative sterol is 25-HC (25-hydroxycholesterol), which can interact with INSIG and induce signalling through SREBP downstream (*17, 49, 50*). Previously, the cryo-EM structures of the INSIG-SCAP complex showed a structure with 25-HC inserted in the dimerising surface, TM3-4 of INSIG, and TM2,4-6 of SCAP (PDB ID: 6M49) (*17*). This leads to a cholesterol glue hypothesis where cholesterol behaves as a stabilising glue between INSIG and SCAP. However, our CG-MD simulation results show that cholesterol binding does not take place at the binding interface nor at the sterol-sensing domain on TM2-6 (Figure 1). However, these regions’ RMSF is affected as a consequence of cholesterol binding in the luminal domain (Figure 4D). This suggested that cholesterol binding must take place elsewhere on SCAP, leading to a large-scale conformational change revealing the INSIG binding site.

Our work suggested that the existing SCAP-INSIG structures with 25-HC are in the activated state, i.e. with cholesterol in the luminal domain binding site. When the cholesterol concentration is high, we suspected that the cholesterol is translocated from the transmembrane to the binding site in the luminal domain through a cavity within the protein, gated by loop 7. This mechanism and path of cholesterol translocation are very similar to NPC1 and other cholesterol-sensing domains (Supplementary Figure 7) (*16, 46–48, 51*). Once the cholesterol is translocated to our identified binding site in the luminal domain, the protein undergoes a conformational change, leading to an opening for the interface for INSIG dimerisation (*18*). This observation agrees with the experimental studies, which highlight that the conformation of SCAP is the key determinant of the dimerisation of SCAP-INSIG (*18, 52*).

The key finding in our studies is the role of sterol; 25-HC behaves very differently to cholesterol in terms of its binding site due to an additional hydroxyl group, which increases the number of hydrogen bonds formed between INSIG and cholesterol (Figure 2). Previous studies have highlighted how different sterol analogues exhibit different binding affinity to SCAP and INSIG, influencing the difference in the sterol sensing mechanism (*49*). Our study highlights the pitfalls of substituting cholesterol to the previously identified site, which might be insufficient to identify the sterol binding site, and thus, a careful approach to the sterol binding pathway must be taken both computationally and experimentally.

One of the critical moments during the preparation of the manuscript is the release of a) the cholesterol parameter for MARTINI3 and b) the open-source AI tools to predict the cholesterol binding site – Chai-1 (*20, 43*). Using the MARTINI3 forcefield, which has scaled interaction, we are still able to capture the same cholesterol binding site in the luminal domain (*20*). We are impressed by the ability of MARTINI3 to capture protein interaction reliably in an interface akin to the recently resolved cryo-EM structure (Figure 1D, Supplementary Figure 2) (*16, 17*). Using Chai-1, we capture cholesterol binding sites both at the lumen and at the transmembrane, showing the putative path for cholesterol translocation (Supplementary figure 8). This highlights the power of recent AI-based computational tools as a method to hypothesise events at the molecular level, integrating with physical-based tools such as molecular dynamics or experimental data (*43*).

To conclude, our study highlighted the importance of loop 7 in recognising and gating cholesterol binding in the luminal domain. Using multiple integrative arrays of computational tools, our work generates novel hypotheses highlighting the mechanism of cholesterol sensors by SCAP in the ER membrane. Understanding this fundamental process’s molecular mechanism could lead to therapeutic design under the cholesterol recognition site. Our computational approach to understanding basic molecular biology will hopefully be the first stepping stones to further experimentally elucidate the mechanism behind cholesterol recognition in the near future.

## Acknowledgements

We thank the member of the Molecular Cell Biology laboratory at Mahidol University for their support and advice throughout the project. We also thank Adisorn Panasawatwong for his support at the start of the project. The MD simulations were performed using resources provided by LANTA High Performance Supercomputer, NSTDA, Thailand (program number: lt200023). The project is funded by the LKY Postdoctoral Fellowship and grants from the Center of Excellence on Environmental Health and Toxicology (EHT), OPS, MHESI and Mahidol University, Bangkok, Thailand.

## Author contributions

C.K. designed, performed research, and wrote and revised the manuscript. P.O. designed research, acquired funding and revised the manuscript. T.P. designed the research, acquired funding, and wrote and revised the manuscript.

---

**Supplementary figure 1.**
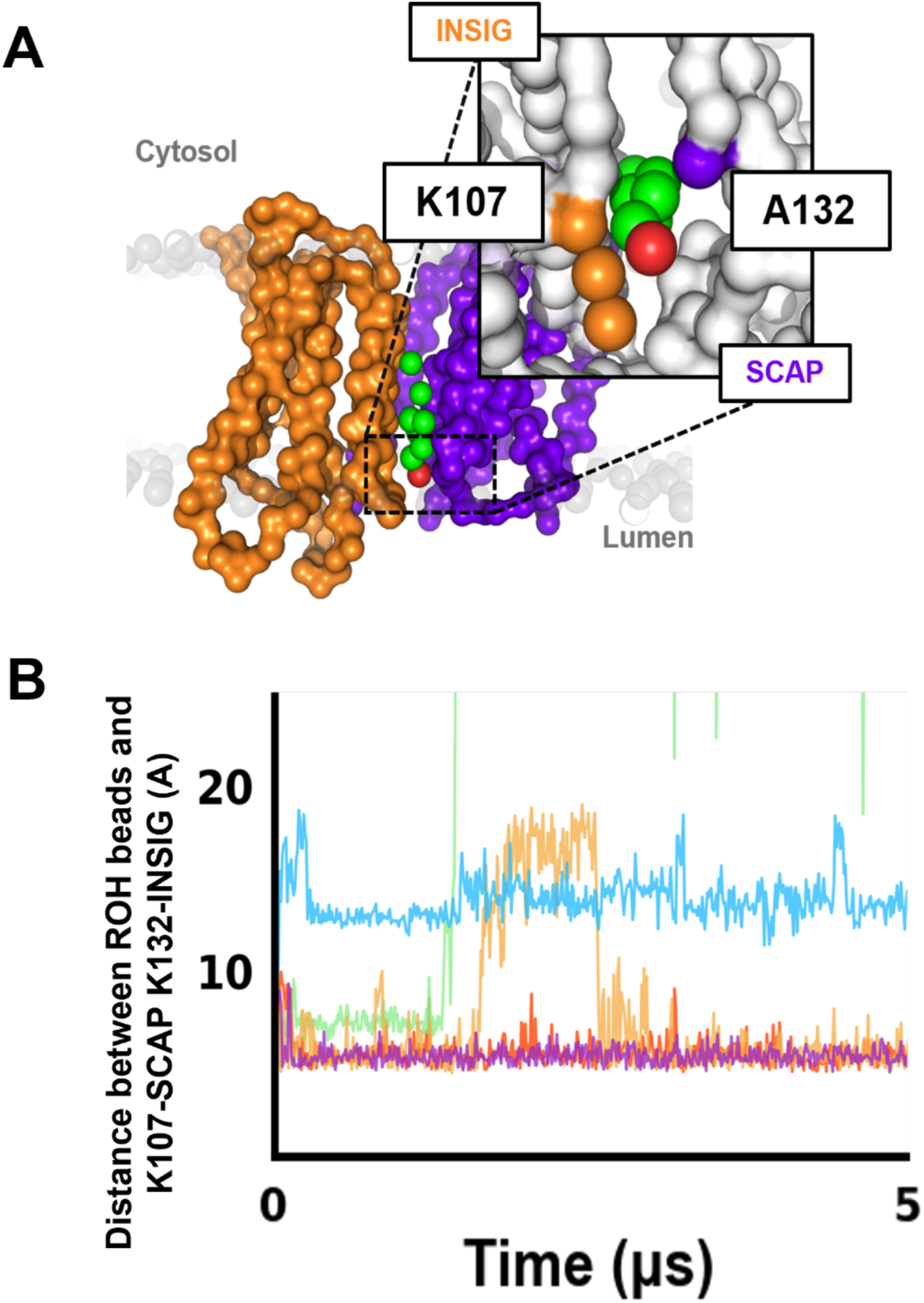
The CG-cholesterol flip-flop within the cleft of TM-complex. **(A)** The CG model of TM-cleft structure shows a cholesterol molecule (green) between TM-INSIG (orange) and TM-SCAP (purple). The reference residues from TM-INSIG and TM-SCAP are depicted in boxes nearby the residues. The ROH head group of cholesterol is shown in red. **(B)** The graph shows changes in the distance between the ROH headgroup of cholesterol and the reference residues from (A). The different colours indicate different repeats (n=5).

**Supplementary figure 2.**
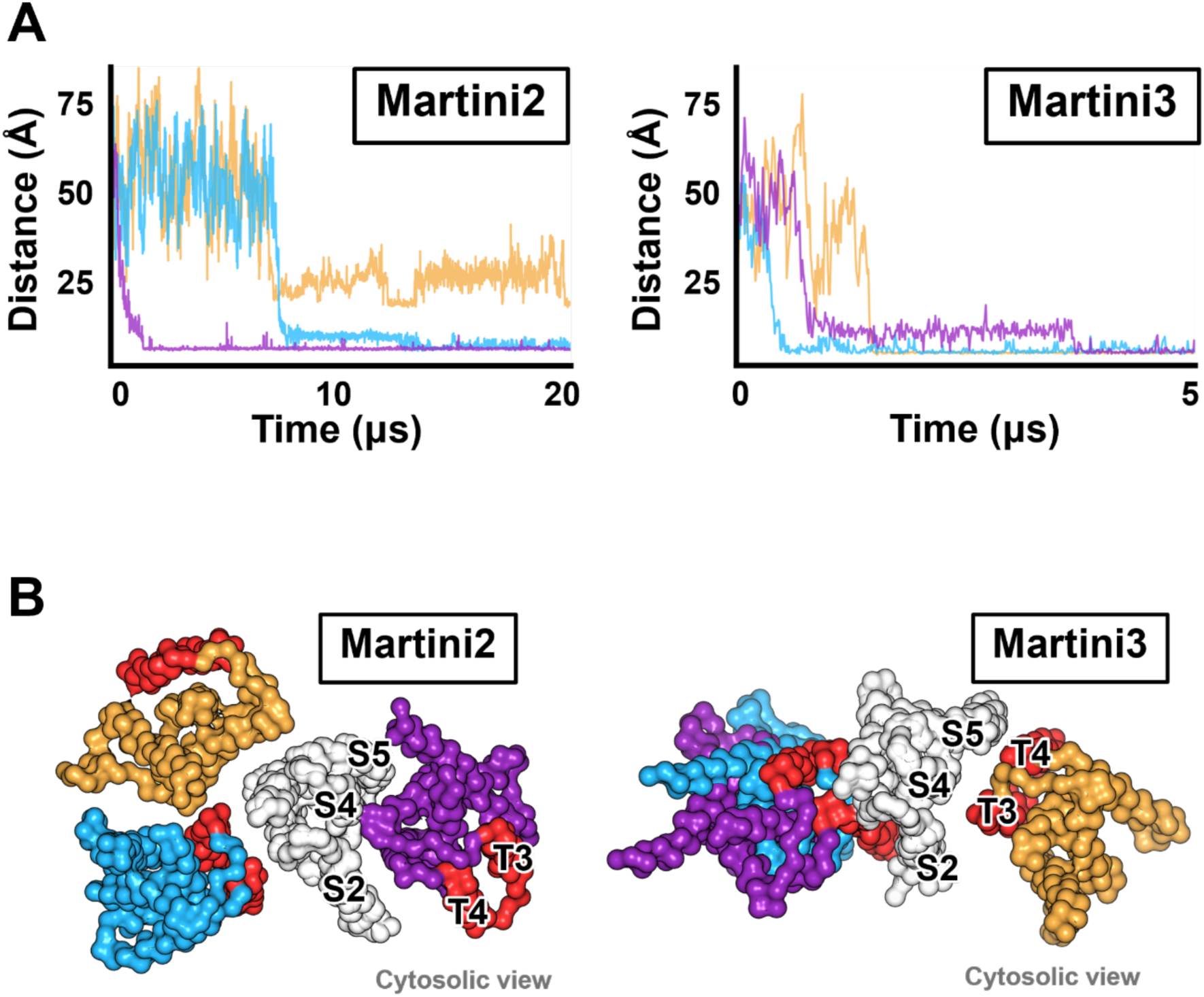
The different results between MARTINI2 and MARTINI3. **(A)** The graphs show differences in distance between TM-INSIG and TM-SCAP in each system using MARTINI2 (left) and Martini3 (right). The different colours indicate different repeats (n=3 of each). **(B)** The top view (cytosolic view) of TM-INSIG and TM-SCAP after 20 µs and 5 µs using MARTINI2 and MARTINI3, respectively, shows different patterns of the dimerisation. The dimerising interface according with the experimental result (PDB ID: 6M49) of TM-INSIG is shown in red (*17*). The secondary structure of the dimerising interface of TM-INSIG and TM-SCAP are labelled with black letters and numbers. The different colours indicate different repeat, as shown in (A).

**Supplementary figure 3.**
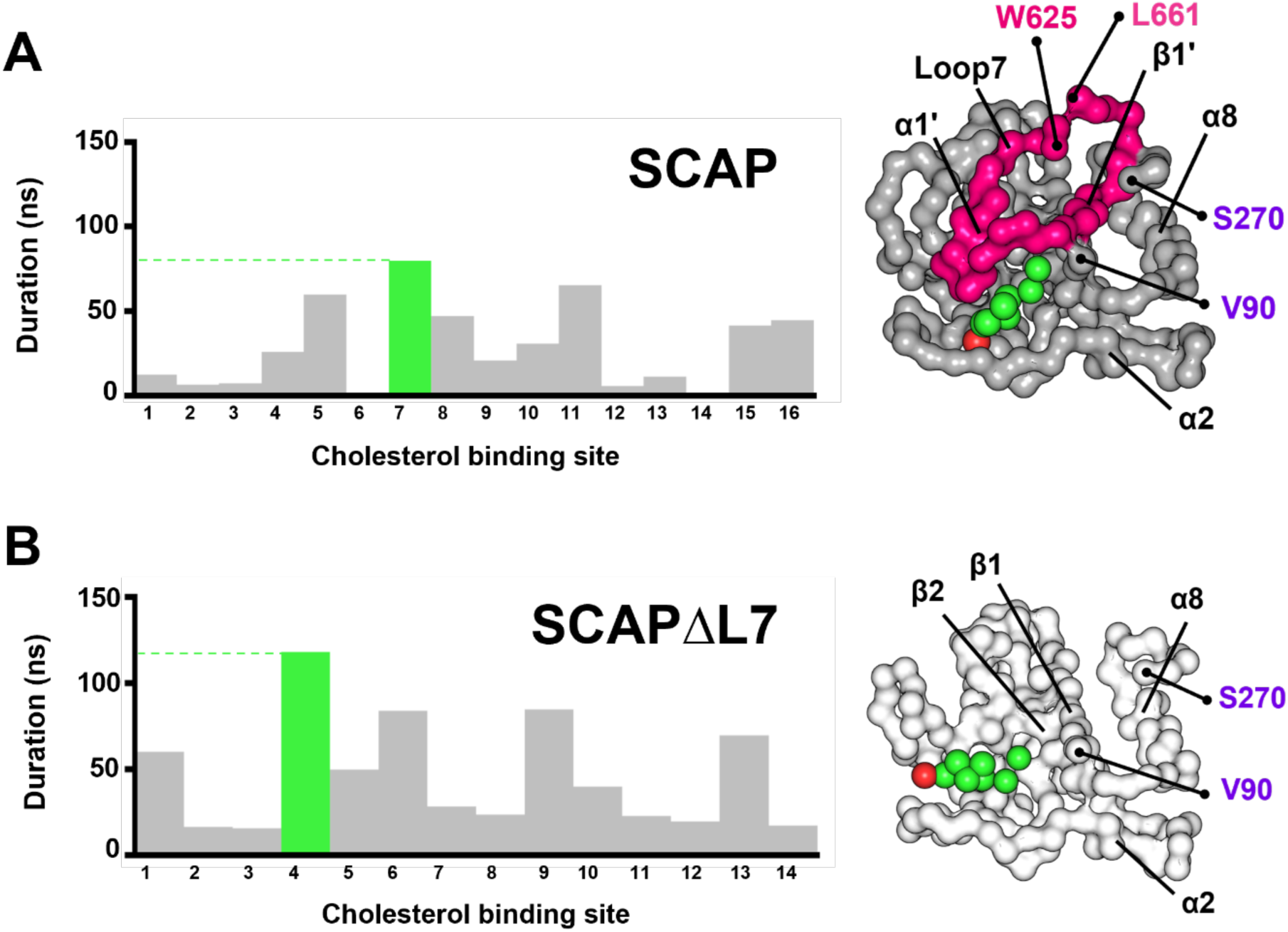
The PyLipID results of SCAP and SCAPΔL7. **(A)-(B)** The comparison of cholesterol binding sites of SCAP (left) and SCAPΔL7 (right) generated by PyLipID shows different durations of cholesterol accession within a dual cut-off (see Material and Methods). The best scores are shown in green and depicted the binding site at right site of each graph. The SCAP system is shown in grey (luminal loop 1) and hot pink (luminal loop 7). The SCAPΔL7 is shown in white. The cholesterol and its ROH head group are shown in green and red, respectively. Each duration scores were calculated from 100 repeats of SCAP and SCAPΔL7 simulations.

**Supplementary figure 4.**
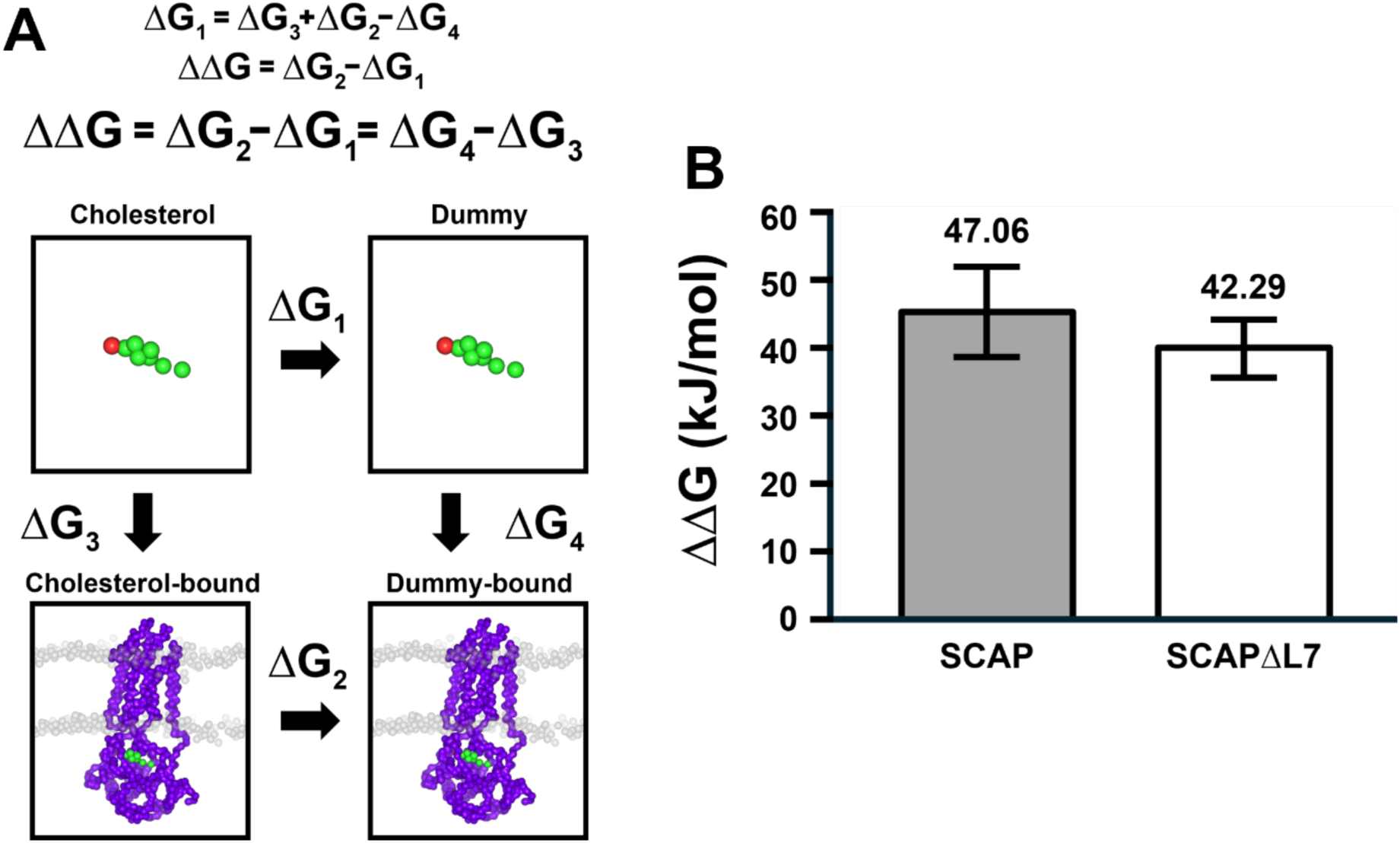
CG-FEP of SCAP and SCAPΔL7. **(A)** The thermodynamic cycle of the CG-FEP shows the calculation of the cholesterol-binding free energy (ΔΔG). The ΔG_1_ is obtained by changing a cholesterol molecule into a dummy in a solution of water and ions. The ΔG_2_ is obtained by the same method but in the binding site on SCAP. The process to obtain ΔG_3_ and ΔG_4_ is the same as applying force to move a cholesterol or a dummy molecule out of the binding site into a solution. **(B)** The CG-FEP results show a difference in cholesterol-binding free energy (ΔΔG) of SCAP (grey) and SCAPΔL7 (white) systems.

**Supplementary figure 5.**
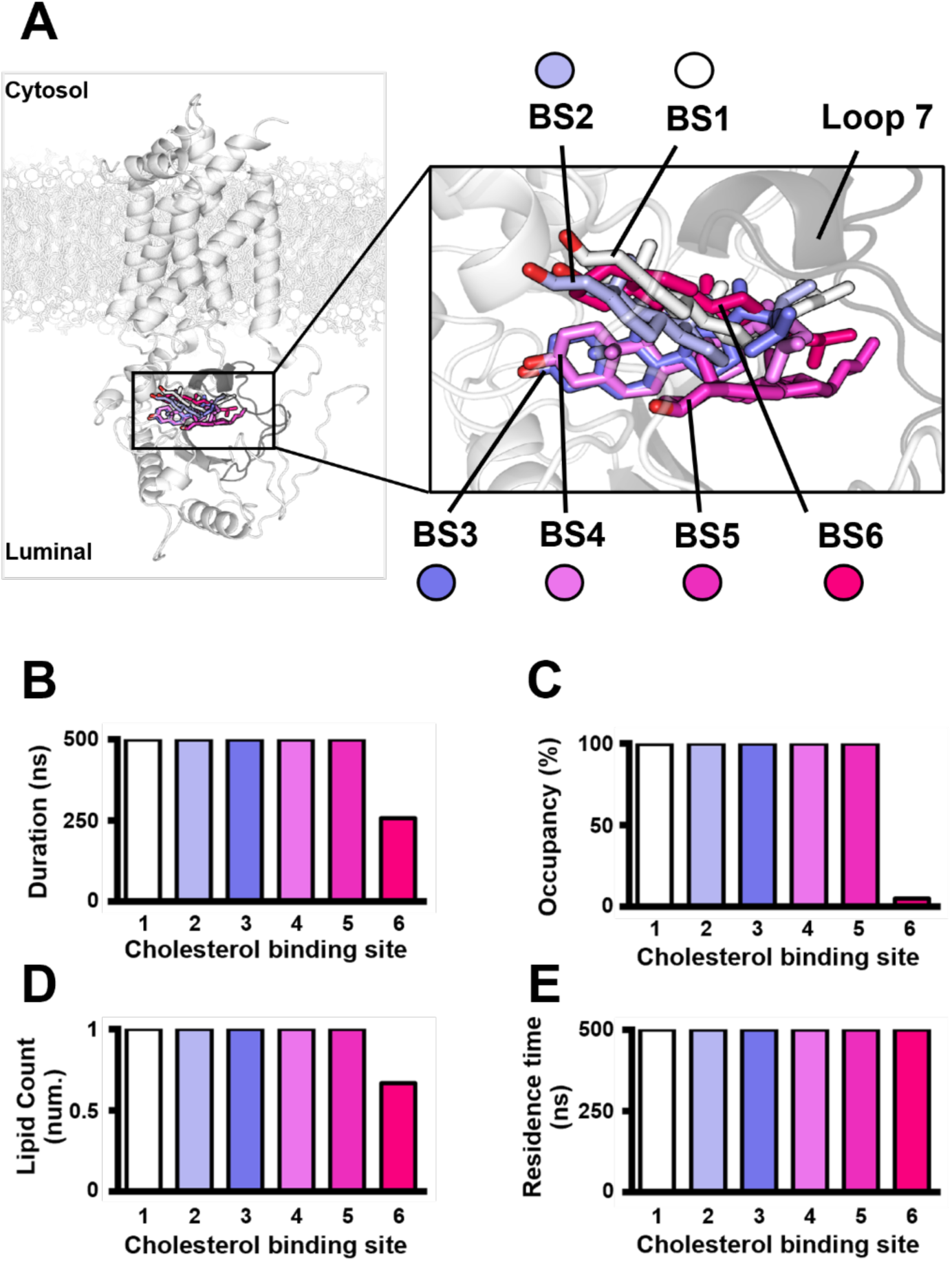
The atomistic detail of cholesterol binding sites from PyLipID. **(A)** The cholesterol binding poses of each binding site generated by PyLipID. The different colours represent a different pose in each cholesterol binding site (BS1-6) within luminal loops 1 (white) and 7 (grey). **(B)-(E)** The data set calculated by PyLipID compares each cholesterol binding site. The different colours represent each cholesterol binding site as shown in (A).

**Supplementary figure 6.**
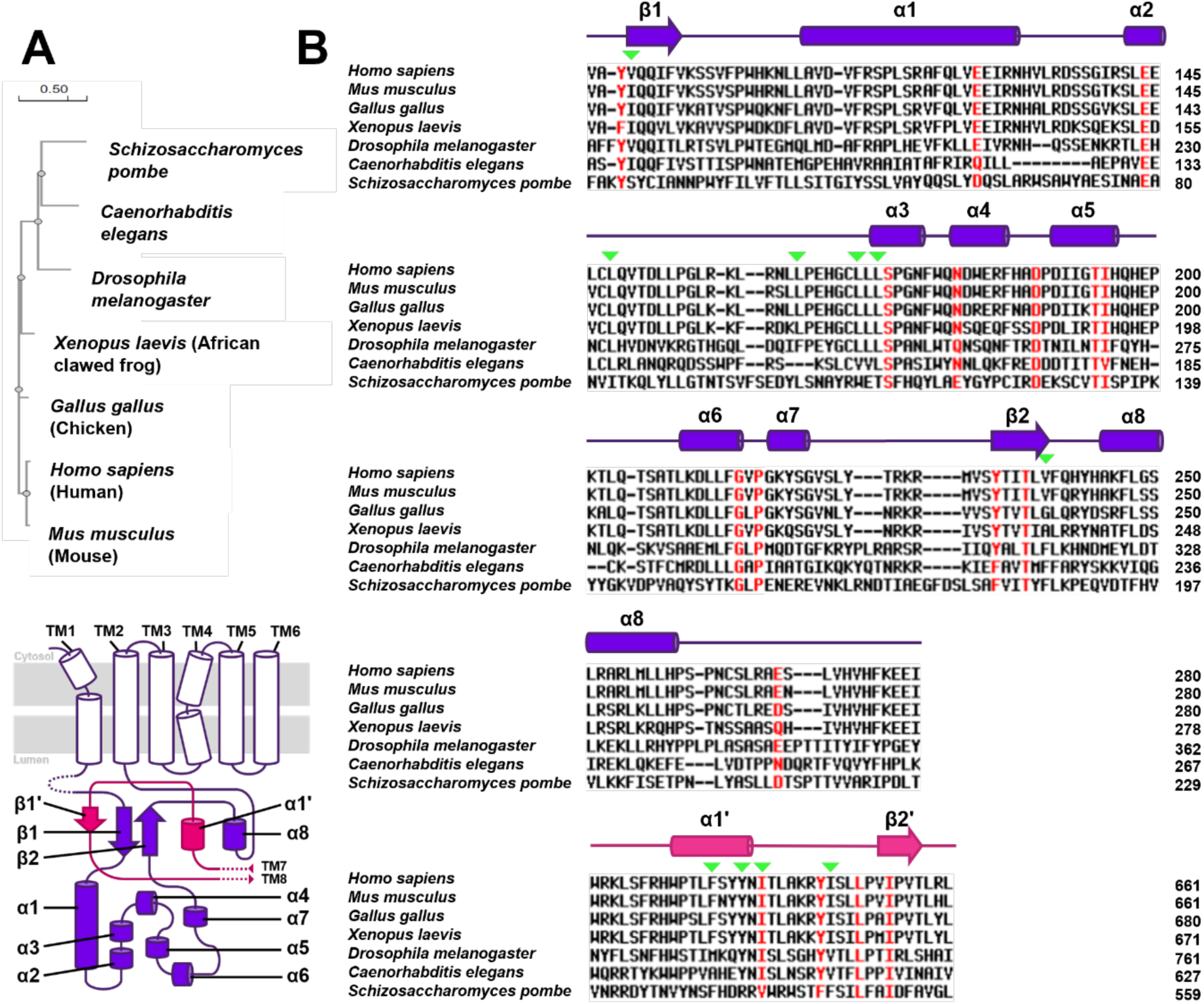
The evolutionary analysis of SCAP in selected organisms. **(A)** The phylogenetic tree of SCAP sequences in selected organisms generated by Clustal Omega. **(B)** The alignments of SCAP sequences generated by the MultAlin web server. The residues that have >95% conservation are shown in red, others are shown in black. The residues of the identified cholesterol binding site are pointed by green marks. The simple topologies of human SCAP luminal loops 1 (residues 90-280) and 7 (residues 625-661) are shown above the alignment corresponding with the sequences. The overall topology of SCAP is shown at the lower left corner of the figure.

**Supplementary figure 7.**
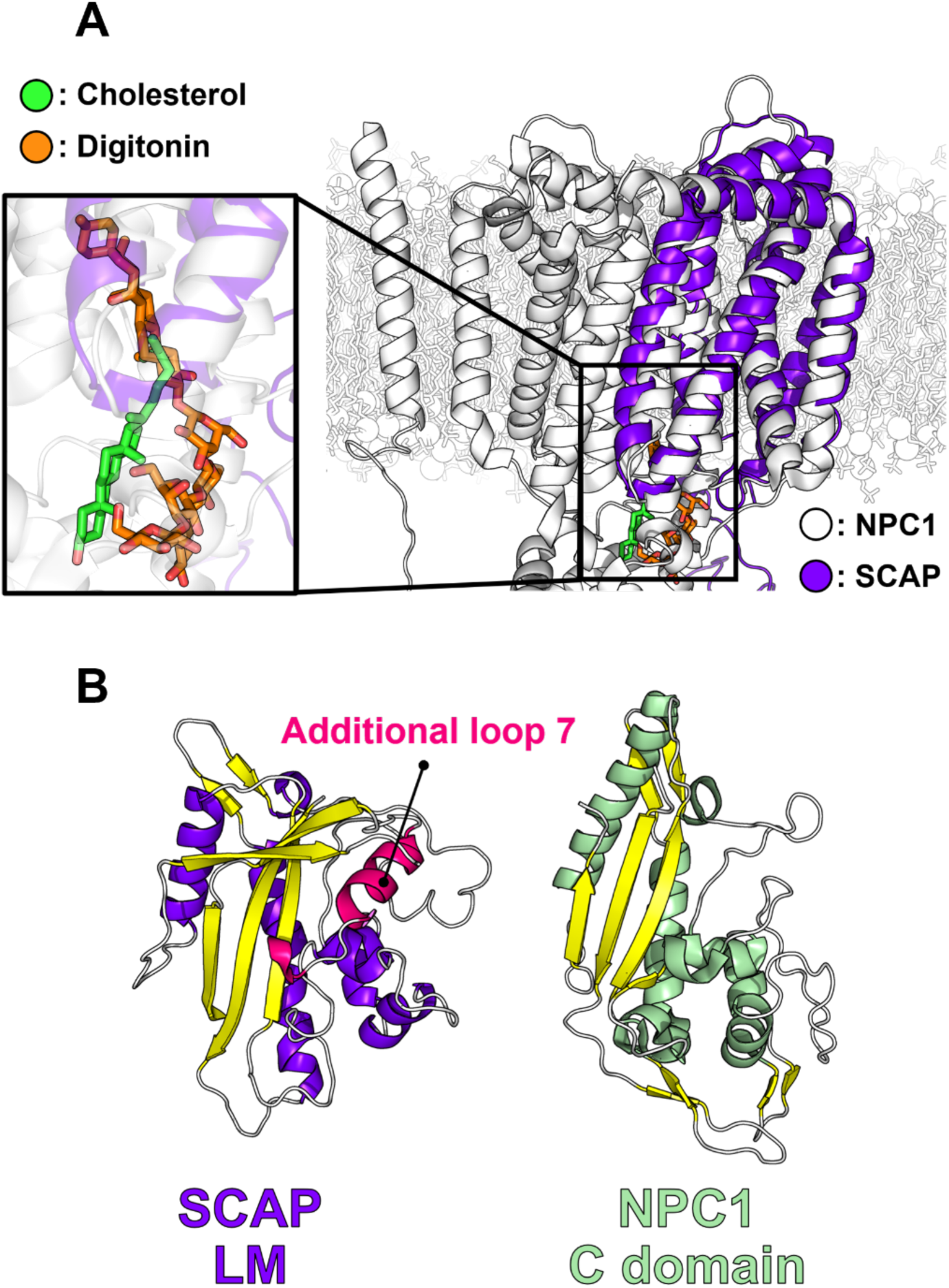
The correlation of SCAP and NPC1. **(A)** The superimposition of SCAP (PDB ID: 7ETW) and NPC1 (PDB ID: 6W5V) shows a similarity between TM domain of SCAP (purple) and NPC1 (white) (*16, 17, 48*). Besides, this reveals an overlapping area of the ligand binding, cholesterol from NPC1 (green) and digitonin from SCAP (orange) (*16, 48*). **(B)** The comparison of the LM domain of SCAP (PDB ID: 7ETW, left) and the C domain of NPC1 (PDB ID: 8EUS, right) shows the similarity of the secondary structures composing in each domain (*16, 47*). The β-sheet is shown in yellow.

**Supplementary figure 8.**
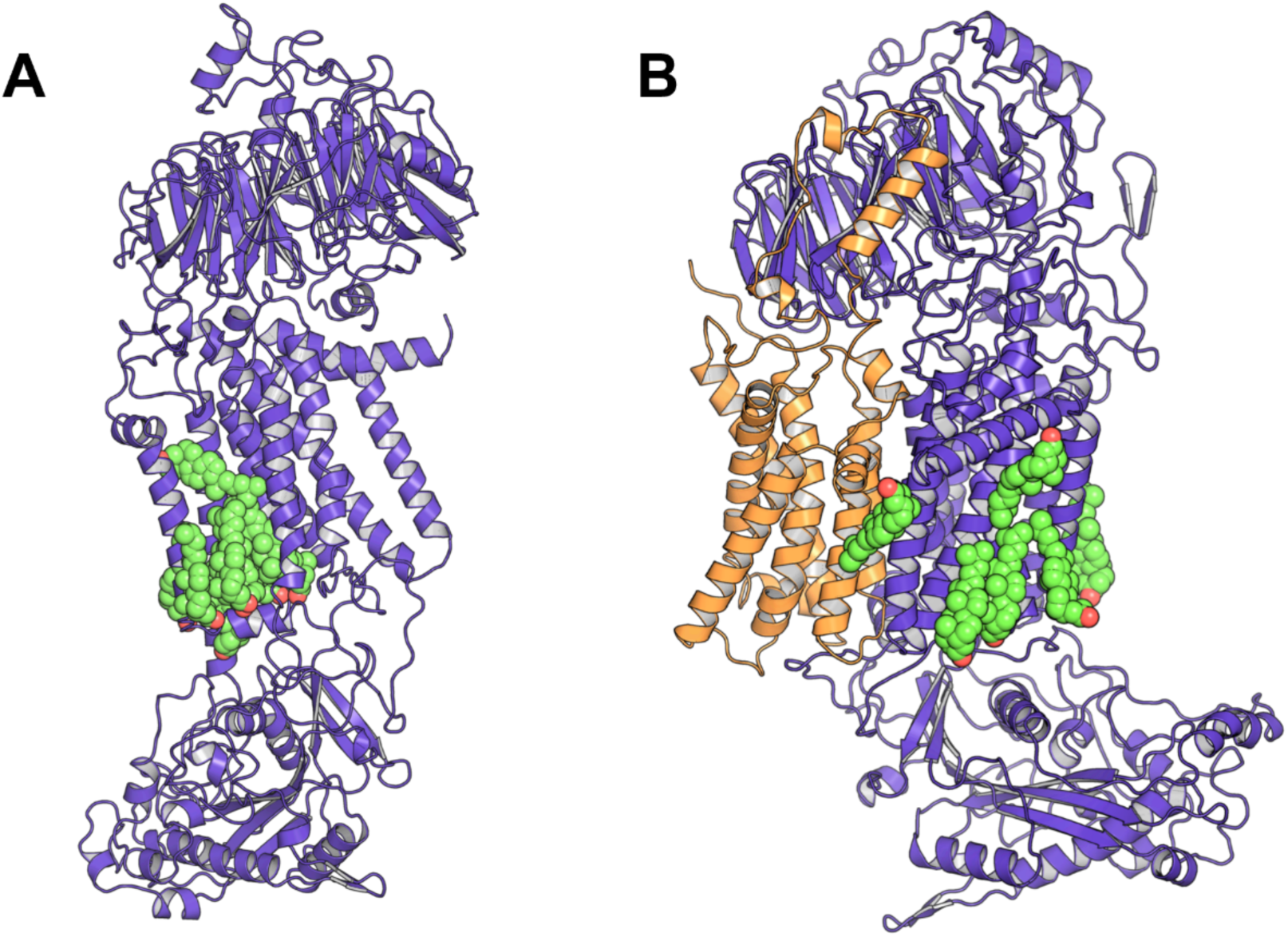
Chai-1 reveals a cluster of cholesterol molecules nearby the LM domain after INSIG dimerisation. The comparison between a full-length SCAP **(A)** and a dimerised complex of full-length INSIG (orange) and SCAP (purple) **(B)** shows a cluster of cholesterol molecules around the SSD domain (TM2/4/5) in (A). After the dimerisation, the cluster of cholesterol molecules disperse around TM1 in (B).

**Supplementary figure 9.**
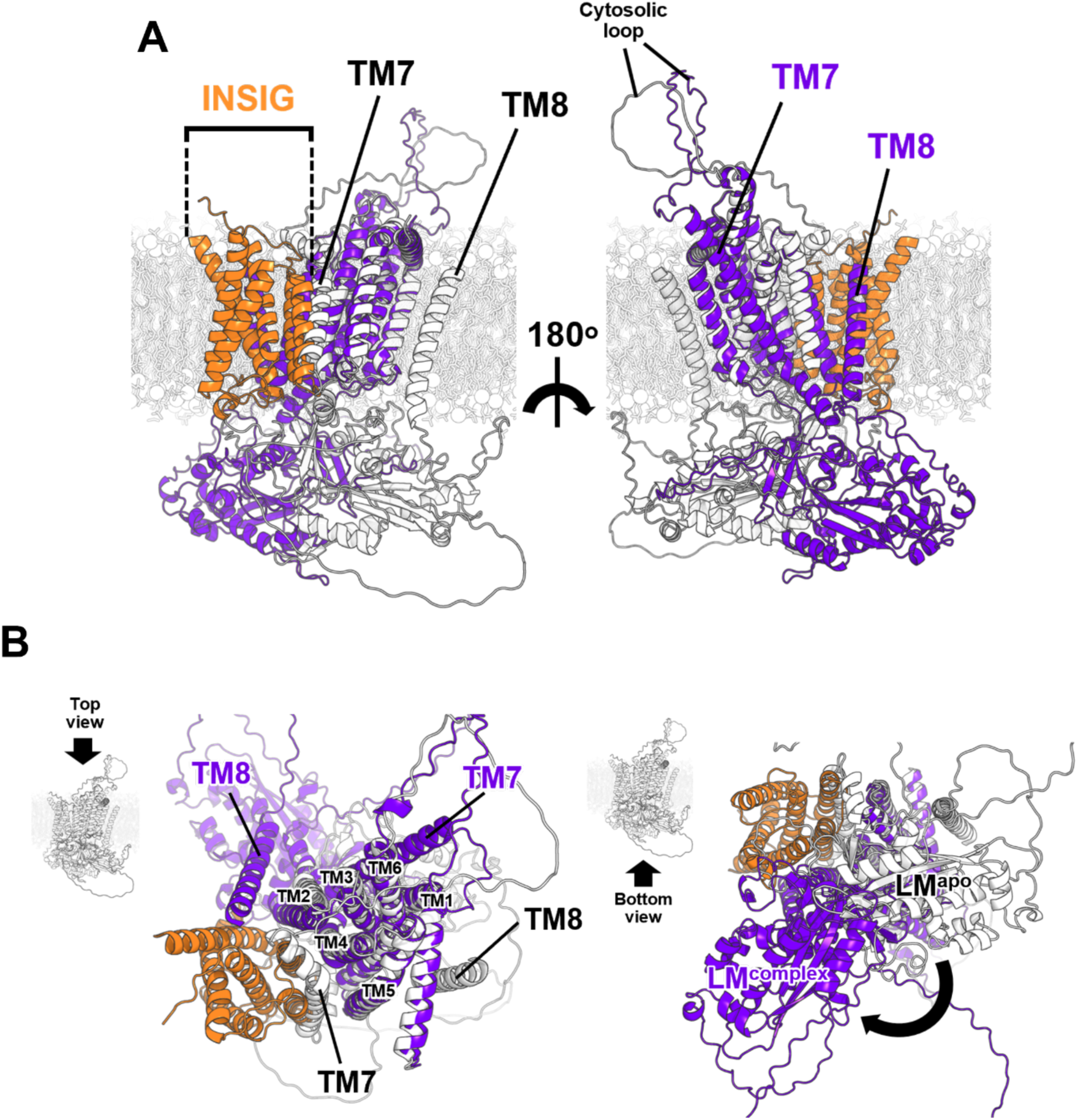
The comparison between SCAP before and after INSIG dimerisation. The superimposition in side-view **(A)**, top-view **(B, left)** and bottom-view **(B, right)** of SCAP (white) and SCAP dimerising with INSIG (purple and orange) shows a distinct movement of the LM domain. Moreover, the superimposition reveals different locations of TM7&8 between before and after INSIG dimerisation. The LM domains of SCAP and SCAP dimerising with INSIG are labelled as LM^apo^ and LM^complex^, respectively.

## References

1. F. R. Maxfield, G. van Meer, Cholesterol, the central lipid of mammalian cells. Curr Opin Cell Biol 22, 422–429 (2010).

2. S. Raffy, J. Teissié, Control of lipid membrane stability by cholesterol content. Biophys J 76, 2072–2080 (1999).

3. T. Harayama, H. Riezman, Understanding the diversity of membrane lipid composition. Nature Reviews Molecular Cell Biology 19, 281–296 (2018).

4. J. Zhang, Q. Li, Y. Wu, D. Wang, L. Xu, Y. Zhang, S. Wang, T. Wang, F. Liu, M. Y. Zaky, S. Hou, S. Liu, K. Zou, H. Lei, L. Zou, Y. Zhang, H. Liu, Cholesterol content in cell membrane maintains surface levels of ErbB2 and confers a therapeutic vulnerability in ErbB2-positive breast cancer. Cell Commun Signal 17, 15 (2019).

5. A. Olżyńska, W. Kulig, H. Mikkolainen, T. Czerniak, P. Jurkiewicz, L. Cwiklik, T. Rog, M. Hof, P. Jungwirth, I. Vattulainen, Tail-Oxidized Cholesterol Enhances Membrane Permeability for Small Solutes. Langmuir 36, 10438–10447 (2020).

6. A. J. Brown, L. J. Sharpe, “Chapter 11 - Cholesterol Synthesis” in Biochemistry of Lipids, Lipoproteins and Membranes (Sixth Edition), N. D. Ridgway, R. S. McLeod, Eds. (Elsevier, Boston, 2016), pp. 327-358.

7. J. Luo, H. Yang, B.-L. Song, Mechanisms and regulation of cholesterol homeostasis. Nature Reviews Molecular Cell Biology 21, 225–245 (2020).

8. J. A. S. Carson, A. H. Lichtenstein, C. A. M. Anderson, L. J. Appel, P. M. Kris-Etherton, K. A. Meyer, K. Petersen, T. Polonsky, L. Van Horn, Dietary Cholesterol and Cardiovascular Risk: A Science Advisory From the American Heart Association. Circulation 141, e39–e53 (2020).

9. I. Tabas, Cholesterol in health and disease. J Clin Invest 110, 583–590 (2002).

10. C. J. Lin, C. K. Lai, M. C. Kao, L. T. Wu, U. G. Lo, L. C. Lin, Y. A. Chen, H. Lin, J. T. Hsieh, C. H. Lai, C. D. Lin, Impact of cholesterol on disease progression. Biomedicine (Taipei) 5, 7 (2015).

11. A. Radhakrishnan, L.-P. Sun, P. J. Espenshade, J. L. Goldstein, M. S. Brown, “Chapter 298 - The SREBP Pathway: Gene Regulation through Sterol Sensing and Gated Protein Trafficking” in Handbook of Cell Signaling (Second Edition), R. A. Bradshaw, E. A. Dennis, Eds. (Academic Press, San Diego, 2010), pp. 2505-2510.

12. A. Radhakrishnan, J. L. Goldstein, J. G. McDonald, M. S. Brown, Switch-like control of SREBP-2 transport triggered by small changes in ER cholesterol: a delicate balance. Cell Metab 8, 512–521 (2008).

13. R. A. DeBose-Boyd, J. Ye, SREBPs in Lipid Metabolism, Insulin Signaling, and Beyond. Trends in Biochemical Sciences 43, 358–368 (2018).

14. A. Radhakrishnan, L. P. Sun, H. J. Kwon, M. S. Brown, J. L. Goldstein, Direct binding of cholesterol to the purified membrane region of SCAP: mechanism for a sterol-sensing domain. Mol Cell 15, 259–268 (2004).

15. L.-P. Sun, J. Seemann, J. L. Goldstein, M. S. Brown, Sterol-regulated transport of SREBPs from endoplasmic reticulum to Golgi: Insig renders sorting signal in Scap inaccessible to COPII proteins. Proceedings of the National Academy of Sciences 104, 6519–6526 (2007).

16. R. Yan, P. Cao, W. Song, Y. Li, T. Wang, H. Qian, C. Yan, N. Yan, Structural basis for sterol sensing by Scap and Insig. Cell Reports 35, (2021).

17. R. Yan, P. Cao, W. Song, H. Qian, X. Du, H. W. Coates, X. Zhao, Y. Li, S. Gao, X. Gong, X. Liu, J. Sui, J. Lei, H. Yang, A. J. Brown, Q. Zhou, C. Yan, N. Yan, A structure of human Scap bound to Insig-2 suggests how their interaction is regulated by sterols. Science 371, eabb2224 (2021).

18. D. L. Kober, A. Radhakrishnan, J. L. Goldstein, M. S. Brown, L. D. Clark, X.-c. Bai, D. M. Rosenbaum, Scap structures highlight key role for rotation of intertwined luminal loops in cholesterol sensing. Cell 184, 3689–3701.e3622 (2021).

19. S. J. Marrink, H. J. Risselada, S. Yefimov, D. P. Tieleman, A. H. de Vries, The MARTINI Force Field: Coarse Grained Model for Biomolecular Simulations. The Journal of Physical Chemistry B 111, 7812–7824 (2007).

20. P. C. T. Souza, R. Alessandri, J. Barnoud, S. Thallmair, I. Faustino, F. Grünewald, I. Patmanidis, H. Abdizadeh, B. M. H. Bruininks, T. A. Wassenaar, P. C. Kroon, J. Melcr, V. Nieto, V. Corradi, H. M. Khan, J. Domański, M. Javanainen, H. Martinez-Seara, N. Reuter, R. B. Best, I. Vattulainen, L. Monticelli, X. Periole, D. P. Tieleman, A. H. de Vries, S. J. Marrink, Martini 3: a general purpose force field for coarse-grained molecular dynamics. Nature Methods 18, 382–388 (2021).

21. S. Yang, C. Song, Switching Go̅-Martini for Investigating Protein Conformational Transitions and Associated Protein–Lipid Interactions. Journal of Chemical Theory and Computation 20, 2618–2629 (2024).

22. K. Mitra, I. Ubarretxena-Belandia, T. Taguchi, G. Warren, D. M. Engelman, Modulation of the bilayer thickness of exocytic pathway membranes by membrane proteins rather than cholesterol. Proc Natl Acad Sci U S A 101, 4083–4088 (2004).

23. T. A. Wassenaar, H. I. Ingólfsson, R. A. Böckmann, D. P. Tieleman, S. J. Marrink, Computational Lipidomics with insane: A Versatile Tool for Generating Custom Membranes for Molecular Simulations. Journal of Chemical Theory and Computation 11, 2144–2155 (2015).

24. Z. Wu, S. Newstead, P. C. Biggin, The KDEL trafficking receptor exploits pH to tune the strength of an unusual short hydrogen bond. Sci Rep 10, 16903 (2020).

25. Schrodinger, LLC. (2015).

26. M. J. Abraham, T. Murtola, R. Schulz, S. Páll, J. C. Smith, B. Hess, E. Lindahl, GROMACS: High performance molecular simulations through multi-level parallelism from laptops to supercomputers. SoftwareX 1, 19–25 (2015).

27. E. J. Haug, J. S. Arora, K. Matsui, A steepest-descent method for optimization of mechanical systems. Journal of Optimization Theory and Applications 19, 401–424 (1976).

28. G. Bussi, D. Donadio, M. Parrinello, Canonical sampling through velocity rescaling. J Chem Phys 126, 014101 (2007).

29. M. Parrinello, A. Rahman, Polymorphic transitions in single crystals: A new molecular dynamics method. Journal of Applied Physics 52, 7182–7190 (1981).

30. W. Song, R. A. Corey, T. B. Ansell, C. K. Cassidy, M. R. Horrell, A. L. Duncan, P. J. Stansfeld, M. S. P. Sansom, PyLipID: A Python Package for Analysis of Protein–Lipid Interactions from Molecular Dynamics Simulations. Journal of Chemical Theory and Computation 18, 1188–1201 (2022).

31. T. Pipatpolkai, R. A. Corey, P. Proks, F. M. Ashcroft, P. J. Stansfeld, Evaluating inositol phospholipid interactions with inward rectifier potassium channels and characterising their role in disease. Communications Chemistry 3, 147 (2020).

32. M. Fajer, R. V. Swift, J. A. McCammon, Using multistate free energy techniques to improve the efficiency of replica exchange accelerated molecular dynamics. J Comput Chem 30, 1719–1725 (2009).

33. P. V. Klimovich, M. R. Shirts, D. L. Mobley, Guidelines for the analysis of free energy calculations. Journal of Computer-Aided Molecular Design 29, 397–411 (2015).

34. E. L. Wu, X. Cheng, S. Jo, H. Rui, K. C. Song, E. M. Dávila-Contreras, Y. Qi, J. Lee, V. Monje-Galvan, R. M. Venable, J. B. Klauda, W. Im, CHARMM-GUI Membrane Builder toward realistic biological membrane simulations. J Comput Chem 35, 1997–2004 (2014).

35. O. N. Vickery, P. J. Stansfeld, CG2AT2: an Enhanced Fragment-Based Approach for Serial Multi-scale Molecular Dynamics Simulations. Journal of Chemical Theory and Computation 17, 6472–6482 (2021).

36. S. Kim, J. Lee, S. Jo, C. L. Brooks III, H. S. Lee, W. Im, CHARMM-GUI ligand reader and modeler for CHARMM force field generation of small molecules. Journal of Computational Chemistry 38, 1879–1886 (2017).

37. J. Huang, A. D. MacKerell, CHARMM36 alŒ atom additive protein force field: Validation based on comparison to NMR data. Journal of Computational Chemistry 34, 2135-2145 (2013).

38. H. J. C. Berendsen, J. P. M. Postma, W. F. v. Gunsteren, A. DiNola, J. R. Haak, Molecular dynamics with coupling to an external bath. The Journal of Chemical Physics 81, 3684–3690 (1984).

39. F. Corpet, Multiple sequence alignment with hierarchical clustering. Nucleic Acids Res 16, 10881–10890 (1988).

40. T. U. Consortium, UniProt: the universal protein knowledgebase in 2021. Nucleic Acids Research 49, D480–D489 (2020).

41. F. Sievers, A. Wilm, D. Dineen, T. J. Gibson, K. Karplus, W. Li, R. Lopez, H. McWilliam, M. Remmert, J. Söding, J. D. Thompson, D. G. Higgins, Fast, scalable generation of high-quality protein multiple sequence alignments using Clustal Omega. Mol Syst Biol 7, 539 (2011).

42. J. Abramson, J. Adler, J. Dunger, R. Evans, T. Green, A. Pritzel, O. Ronneberger, L. Willmore, A. J. Ballard, J. Bambrick, S. W. Bodenstein, D. A. Evans, C.-C. Hung, M. O’Neill, D. Reiman, K. Tunyasuvunakool, Z. Wu, A. Žemgulytė, E. Arvaniti, C. Beattie, O. Bertolli, A. Bridgland, A. Cherepanov, M. Congreve, A. I. Cowen-Rivers, A. Cowie, M. Figurnov, F. B. Fuchs, H. Gladman, R. Jain, Y. A. Khan, C. M. R. Low, K. Perlin, A. Potapenko, P. Savy, S. Singh, A. Stecula, A. Thillaisundaram, C. Tong, S. Yakneen, E. D. Zhong, M. Zielinski, A. Žídek, V. Bapst, P. Kohli, M. Jaderberg, D. Hassabis, J. M. Jumper, Accurate structure prediction of biomolecular interactions with AlphaFold 3. Nature 630, 493–500 (2024).

43. J. Boitreaud, J. Dent, M. McPartlon, J. Meier, V. Reis, A. Rogozhonikov, K. Wu, Chai-1: Decoding the molecular interactions of life. bioRxiv, (2024).

44. M. Motamed, Y. Zhang, M. L. Wang, J. Seemann, H. J. Kwon, J. L. Goldstein, M. S. Brown, Identification of luminal Loop 1 of Scap protein as the sterol sensor that maintains cholesterol homeostasis. J Biol Chem 286, 18002–18012 (2011).

45. A. Meller, M. Ward, J. Borowsky, M. Kshirsagar, J. M. Lotthammer, F. Oviedo, J. L. Ferres, G. R. Bowman, Predicting locations of cryptic pockets from single protein structures using the PocketMiner graph neural network. Nature Communications 14, 1177 (2023).

46. N. Elghobashi-Meinhardt, Cholesterol Transport in Wild-Type NPC1 and P691S: Molecular Dynamics Simulations Reveal Changes in Dynamical Behavior. International Journal of Molecular Sciences 21, 2962 (2020).

47. L. Odongo, K. K. Zadrozny, W. E. Diehl, J. Luban, J. M. White, B. K. Ganser-Pornillos, L. K. Tamm, O. Pornillos, Purification and structure of luminal domain C of human Niemann-Pick C1 protein. Acta Crystallogr F Struct Biol Commun 79, 45–50 (2023).

48. H. Qian, X. Wu, X. Du, X. Yao, X. Zhao, J. Lee, H. Yang, N. Yan, Structural Basis of Low-pH-Dependent Lysosomal Cholesterol Egress by NPC1 and NPC2. Cell 182, 98–111.e118 (2020).

49. A. Radhakrishnan, Y. Ikeda, H. J. Kwon, M. S. Brown, J. L. Goldstein, Sterol-regulated transport of SREBPs from endoplasmic reticulum to Golgi: oxysterols block transport by binding to Insig. Proc Natl Acad Sci U S A 104, 6511–6518 (2007).

50. C. M. Adams, J. Reitz, J. K. De Brabander, J. D. Feramisco, L. Li, M. S. Brown, J. L. Goldstein, Cholesterol and 25-Hydroxycholesterol Inhibit Activation of SREBPs by Different Mechanisms, Both Involving SCAP and Insigs*. Journal of Biological Chemistry 279, 52772–52780 (2004).

51. T. Long, X. Qi, A. Hassan, Q. Liang, J. K. De Brabander, X. Li, Structural basis for itraconazole-mediated NPC1 inhibition. Nature Communications 11, 152 (2020).

52. Y. Gao, Y. Zhou, J. L. Goldstein, M. S. Brown, A. Radhakrishnan, Cholesterol-induced conformational changes in the sterol-sensing domain of the Scap protein suggest feedback mechanism to control cholesterol synthesis. Journal of Biological Chemistry 292, 8729–8737 (2017).

